# Intact and single-molecule analysis of heparan sulfate

**DOI:** 10.64898/2026.06.26.734651

**Authors:** Petar Hristov, Pooria Daneshvar Kakhaki, Talia Tzadikario, Sandeep K. Rai, Guowei Su, Paulo H. Olivieri, Jeffrey D. Esko, Jian Liu, Miten Jain, Ryan A. Flynn

## Abstract

Establishing tools to couple biological processes to a DNA sequence has transformed our ability to monitor life at the molecular scale due to the scalability, flexibility, and low cost of DNA sequencing. Key examples include DNA-protein (ChIP-seq^1^), RNA-protein (CLIP-seq^2^), protein-protein (proximity ligation assay^3^), and Cas-based recording of cellular events^4^. In contrast, this paradigm has not yet significantly enhanced studies of glycans, which are mostly limited to non-DNA based chemical and biochemical assays. While classical asparagine-linked and serine/threonine-linked glycans can be directly sequenced using mass spectrometry, glycosaminoglycans - notable players in the extracellular matrix - cannot be easily analyzed in their full-length form. Here we introduce HS-nano-seq, a generalized framework to selectively label, process, and detect features of heparan sulfate on a nanopore sequencing platform. Recognizing that heparan sulfate is biochemically analogous to a nucleic acid, we report purification techniques using rapid nucleic acid strategies and conjugation methods to couple DNA adapters, generating HS-DNA chimeras resolved as discrete species by capillary electrophoresis (CE). The CE assay can distinguish features of chain length and sulfation patterns. At the single-molecule level enabled by nanopore sensing, we classify a library of synthetic heparan sulfate standards and demonstrate that nanopore ionic current fingerprints encode sulfation-dependent structural features of individual HS chains. Analysis of intact, cell-derived HS could discriminate features of individual chains with different sulfation patterns, defining the heterogeneity of binding motifs across cell types and how cells organize and program the tethered extracellular matrix. More broadly, HS-nano-seq establishes a framework for achieving full-length readouts of ECM glycopolymers that are amenable to the same biological interrogation as nucleic acids.

## Introduction

The cell surface and extracellular matrix display a rich repertoire of glycans that regulate signaling, adhesion, and molecular recognition^5^. While branched asparagine *N*- and serine/threonine *O*-linked glycans can be structurally profiled by mass spectrometry^6^, intact glycosaminoglycan chains (GAGs) remain difficult to analyze^7^. GAGs are linear, highly anionic polysaccharides that include heparan sulfate (HS), heparin (HP), chondroitin sulfate (CS), dermatan sulfate (DS), keratan sulfate (KS), and hyaluronic acid (HA)^8^. Heparan sulfate, in particular, is broadly expressed on cell surfaces and in the extracellular matrix (ECM). HS is synthesized as a linear polysaccharide composed of alternating D-glucuronic acid (GlcA) and *N*-acetyl-D-glucosamine (GlcNAc) residues, polymerized in the Golgi apparatus by the heterodimeric exostosin complex EXT1/2. This nascent backbone is subsequently diversified by a stepwise enzymatic cascade: *N*-deacetylase/*N*-sulfotransferases (NDST1-4) convert select GlcNAc residues into *N*-sulfated glucosamine; HS C5-epimerase converts adjacent D-GlcA to L-iduronic acid (IdoA), HS2ST can introduce sulfate groups at C2 position of the uronic acids; and 6-*O*-sulfotransferases (HS6ST1-3) install 6-*O*-sulfates on select GlcNAc and GlcNS residues. Additional structural diversity arises from rare but functionally potent 3-*O*-sulfation events catalyzed by HS3ST1-6. The assembly process is non-template driven and includes post-assembly processing by cell surface endo-glucosaminidase (HPSE) and endo-6-sulfatases (SULFs). The arrangement of these sulfated domains and the degree of sulfation generate highly heterogeneous sulfation patterns that remain difficult to interrogate in a full-length sequence context.

Importantly, these sulfation patterns determine how HS engages its binding partners. Sulfation domains along HS chains serve as interaction sites for a wide range of extracellular proteins, including growth factors, apolipoproteins, proteinase inhibitors, cytokines, viral particles, and cell surface glycoRNAs^9,10,11^. Binding is driven by the arrangement and composition of sulfated, iduronic acid-rich regions known as S-domains^12^. The most well-characterized GAG-protein interaction is the 3-*O*-sulfated pentasaccharide motif in heparin - a highly sulfated form of HS - which binds antithrombin III (ATIII) with high specificity and endows heparin with anticoagulant activity^13^. Defined synthetic HS oligosaccharides have revealed preferences for different sulfation motifs for other proteins such as growth factors, including members of the fibroblast growth factor (FGF) and vascular endothelial growth factor (VEGF) families of proteins^14^, but the lack of sequencing methods has limited the ability to build structure-activity relationships^15^.

A major barrier to a sulfation domain-centric understanding of the ECM is the lack of analytical methods that can read glycosaminoglycans in their intact, full-length form. Although mass spectrometry is the principal tool for glycan structural analysis, native heparan sulfate chains (and other GAGs) are particularly challenging because chain-length heterogeneity, variable sulfation and epimerization, and multiple charge-state distributions generate highly complex spectra^7^. As a result, full-length GAG sequencing by MS-based top-down workflows has been largely restricted to select simple systems, such as the single, short, and more-or-less homogeneous CS/DS chains found on the proteoglycans bikunin and decorin^16,17^. Typical HS analysis instead relies on bottom-up MS after enzymatic or chemical depolymerization, which has revealed tissue- and cell-type-specific disaccharide sulfation profiles^18^, but erases the positional arrangement of those disaccharides along individual chains. Thus, the sequence-level organization of HS sulfation domains remains largely inaccessible.

As a linear, highly anionic polymer, full-length heparan sulfate resembles a nucleic acid in both architecture and electrostatics. This analogy has motivated us and others^19,20,21^ to explore nanopore-based strategies for intact HS analysis. Previous studies have established two key foundations for this approach: GAGs can translocate through biological nanopores^19^, and differences in sulfation can measurably alter ionic-current signatures^20,21^. Yet, intact chains move through pores rapidly and stochastically, limiting monomer-resolved readout along the polymer^22,23^. In DNA nanopore sequencing, single-nucleotide resolution became possible only when motor proteins were introduced to ratchet strands through the pore in slow, discrete steps^24^. We therefore asked whether analogous motor-controlled translocation of HS could convert nanopore sensing into a spatially resolved readout of sulfation-domain organization.

Here, we report the optimization of DNA purification, processing, and sensing technologies for the purpose of characterizing GAGs with a focus on HS. These tools allow for rapid handling of intact and native HS chains with characterization of the lengths and abundance in bulk using standard and capillary electrophoresis systems. By creating DNA-HS-DNA chimeras, we enabled nanopore interrogation of synthetic and native, cell-derived HS chains, allowing the direct observation of molecular features at the single-molecule level.

## Results

### Heparan sulfate can be rapidly purified and mono-labeled at both ends of the chain

To analyze intact heparan sulfate chains by nanopore sensing, we first selectively derivatized both the reducing end (RE) and the non-reducing end (NRE) of the chains to enable end-to-end analysis (Fig. 1a). Reductive amination at the RE is widely used to install UV- and fluorogenic tags because of the reactive carbonyl group on the open ring form of the terminal sugar^25^; however, this reaction is not inherently specific to HS and can label other glycans in complex biological samples after the chains are released by β-elimination or enzymatically. In contrast, the NRE provides a biochemical handle with greater HS selectivity. EXT1/2 which is involved in endogenous chain polymerization can act *in vitro* by adding a single monosaccharide from a UDP-sugar donor^26^. Combining enzymatic NRE labeling with chemical RE labeling therefore provides a route to enrich stringently for intact HS chains bearing both terminal labels.

**Figure 1.**
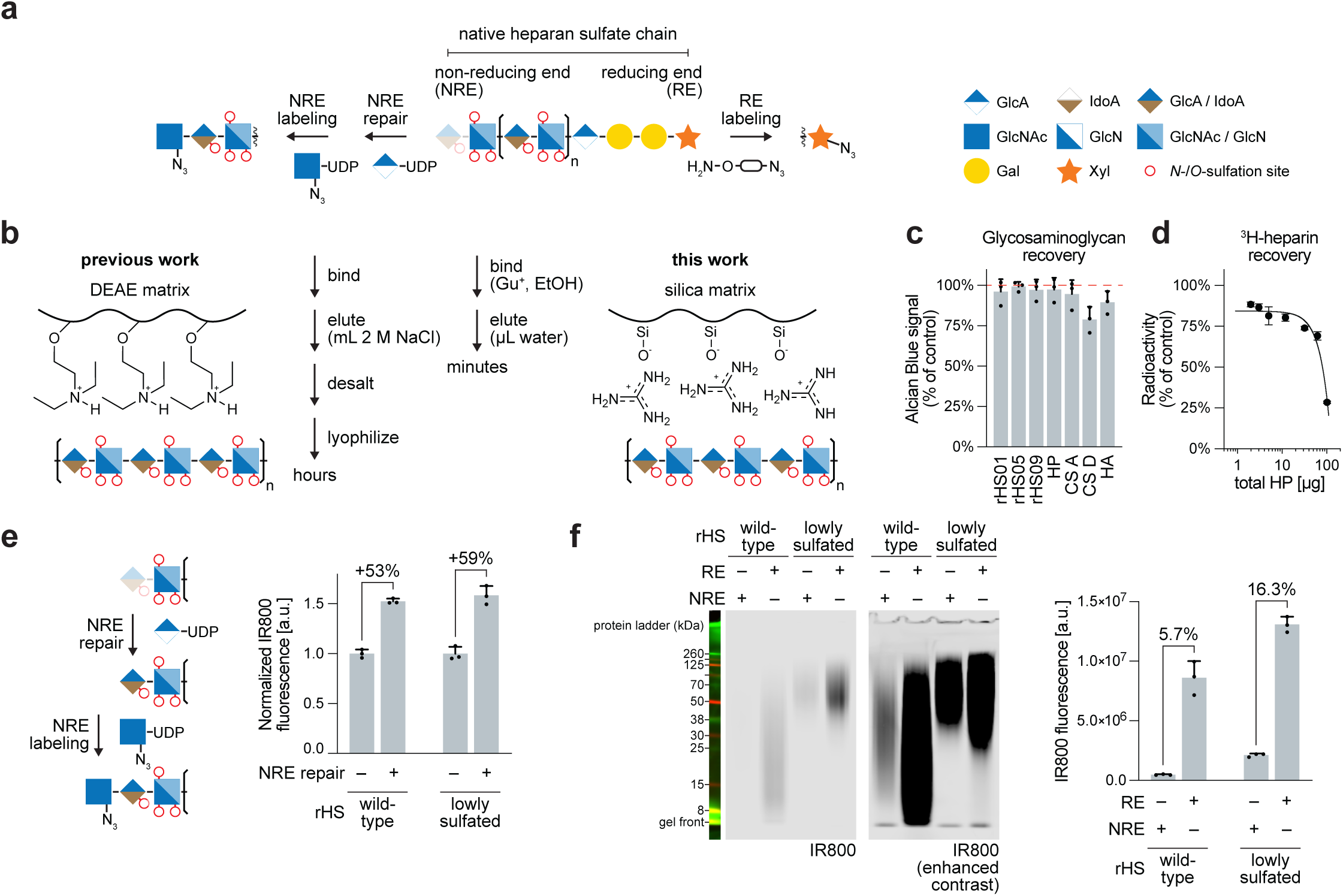
Rapid purification and end-selective mono-labeling of intact heparan sulfate chains. a. Schematic of end-selective mono-labeling of intact heparan sulfate (HS). PmHS2 installs a single azide-modified GlcNAc from UDP-GlcNAz onto GlcA-capped non-reducing ends (NREs); GlcNAc-capped NREs are first repaired by PmHS2-mediated addition of GlcA from UDP-GlcA. The reducing end (RE) is chemically functionalized by hydroxylamination using an aminooxy-azide linker. Sugar symbols follow SNFG notation^45,46^; red circles indicate potential sulfation sites. b. Conventional DEAE anion-exchange purification requires high-salt elution, desalting, and lyophilization. Silica column-based purification of HS uses guanidinium-mediated binding followed by rapid elution in water. c. Recovery of representative glycosaminoglycans after silica purification, quantified by Alcian Blue staining. Analytes include cell-derived HS of varying sulfation levels (rHS01, rHS05, rHS09), heparin (HP), chondroitin sulfate (CS-A, CS-D), and hyaluronic acid (HA). Data from Supplementary Fig. 1a. n = 3; bars indicate mean ± s.d. d. Recovery of [^3^H]heparin after silica purification. [^3^H]heparin was spiked into increasing amounts of unlabeled heparin, purified over a silica column, and eluate fractions were quantified by liquid scintillation counting. Curve represents a nonlinear regression fit to a one-site total binding model (GraphPad Prism). n = 2; bars indicate mean ± s.d. e. Wild-type (rHS01) and lowly-sulfated (rHS05) HS treated with UDP-GlcA with (+) or without (-) PmHS2 before UDP-GlcNAz labeling and IR800 detection. Data from Supplementary Fig. 2f. n = 3; bars indicate mean ± s.d. Percentages indicate the increase in IR800 fluorescence after NRE repair. f. Comparison of end-selective labeling of rHS01 and rHS05 assessed by SDS-PAGE with IR800 detection. HS chains were labeled enzymatically at the NRE or chemically at the RE. Right, quantification of IR800 fluorescence from labelled HS species. Data from Supplementary Fig. 3f. n = 3; bars indicate mean ± s.d. Percentages indicate NRE labeling signal relative to the corresponding RE-labeled sample.

Repeated chemical and enzymatic manipulation of intact HS chains requires a rapid purification method compatible with iterative cloning workflows. Conventional HS purification relies on anion-exchange chromatography followed by high-salt elution (1-2 M NaCl), desalting, and lyophilization - a lengthy process that takes 1-3 days^27^. Given the negative charge of the HS chains, we reasoned that silica-based purification routinely used for nucleic acids might provide a simpler alternative, allowing direct elution in water within minutes (Fig. 1b). Using electrophoresis and Alcian Blue staining, we found that representative full-length GAGs bind robustly to silica columns and elute with high recovery (Fig. 1c and Figure S1a). Titration of [^3^H]heparin with non-radioactive material allowed determination of binding capacity, which was ≥80% up to 65 µg, and comparable to that of nucleic acid (Fig. 1d). All of the recovered material was susceptible to heparin lyases I, II, and III. The lyases were not retained on the column, enabling sequential enzymatic treatment and purification without additional cleanup (Figure S1b,c). While native HS chains (>100 monosaccharides) were efficiently retained, a synthetic 18-mer was not, indicating a length dependence for silica binding similar to that of nucleic acids (Figure S1d). This one-step, ∼15-minute purification enables rapid iteration of end-labeling reactions on intact HS from biological samples.

Next, we optimized selective enzymatic NRE labeling of intact HS chains. EXT1/2-mediated mono-labeling of GlcA-capped NREs using UDP-GlcNAz has been previously demonstrated^26,28^. We additionally evaluated *Pasteurella multocida* heparosan synthase 2 (PmHS2), a bacterially expressed, single-subunit bifunctional glycosyltransferase widely used in synthetic HS oligosaccharide production^29,30^. PmHS2 also incorporated UDP-GlcNAz into full-length HS chains (Figure S2b), dependent on concurrent presence of HS, PmHS2, and UDP-GlcNAz (Figure S2c). The modified HS remained sensitive to heparin lyases digestion (Figure S2d). Labeling of GlcA-capped NREs reached saturation within ∼30 min (Figure S2e), likely reflecting the high enzyme-to-substrate ratio *in vitro*. Given its ease of production and demonstrated activity on sulfated substrates, we used PmHS2 for all subsequent NRE labeling reactions.

Natural HS chains have heterogeneous NREs, terminating in either GlcA or GlcNAc residues, with additional variability arising from sulfation near the terminus^31,32^. To expand NRE mono-labeling beyond GlcA-capped termini, we implemented an “NRE repair” strategy in which GlcNAc-terminated ends are extended with UDP-GlcA, converting them into GlcA-capped NREs competent for subsequent UDP-GlcNAz mono-labeling. The application of NRE repair to naturally sulfated (rHS01) and minimally sulfated (rHS05) cell-derived HS chains increased the NRE labeling efficiency by approximately 50% (Fig. 1e and Figure S2f), broadening the fraction of native HS chains accessible to NRE-specific derivatization.

Labeling of the reducing end requires release of the HS chains from their heparan sulfate proteoglycan (HSPG) core proteins via β-elimination, generating aldehyde-reactive hemiacetals including truncated “peeling” by-products^33^ (Figure S3a). During disaccharide analysis, this functionality is derivatized by reductive amination with aniline or fluorophores to enable UV, fluorescence or MS-based detection^25,34^ (Figure S3b). As a reducing-agent-free alternative, we used hydroxylamination with aminooxy-PEG_3_-azide to form a stable oxime adduct that installs an azide handle identical in downstream reactivity to the NRE label^35^ (Figure S3c,d). RE labeling reached saturation within 1 h and did not interfere with HSases digestion (Figure S3e,f). As hydroxylamination targets all β-eliminated REs regardless of backbone structure, RE labeling provides an estimate of the total molar mass of HS chains that can be labeled in a sample.

Together, these NRE and RE modifications enabled mono-labeling of either terminus of a native HS chain within hours. Whereas RE labeling captures the total pool of chains, NRE labeling is chemoenzymatic and may therefore be sensitive to the structural context of the terminus. Comparison of the NRE and RE labeling of wild type (rHS01) and lowly-sulfated (rHS05) HS chains revealed that NRE labeling captured only a fraction of the RE signal −5.7% for rHS01 and 16.3% for rHS05 (Fig. 1f and Figure S3g). The higher NRE/RE ratio in the less sulfated species suggests that the observed PmHS2 activity is influenced by sulfation near the NRE.

### HS-DNA conjugates can be sensed with standard capillary electrophoresis

Native heparan sulfate exists as a distribution of chain lengths with variable sulfation patterns, yet no tools exist to profile intact chains at the sensitivity and throughput available for nucleic acids. We reasoned that covalent conjugation of HS to DNA would bridge the gap, reframing HS analysis into a nucleic acid dimension (Fig. 2a). As a proof of concept, we conjugated a non-sulfated synthetic HS 9-mer to a double-stranded DNA adapter and analyzed the product by TapeStation capillary electrophoresis. The HS-DNA conjugate migrated at a higher apparent nucleic acid length than the unconjugated adapter, enabling concentration-dependent monitoring of the conjugate formation (Fig. 2b). TapeStation detection of the HS-DNA conjugate approached the sensitivity of standard nucleic acid measurements (Fig. 2c and Figure S4a).

**Figure 2.**
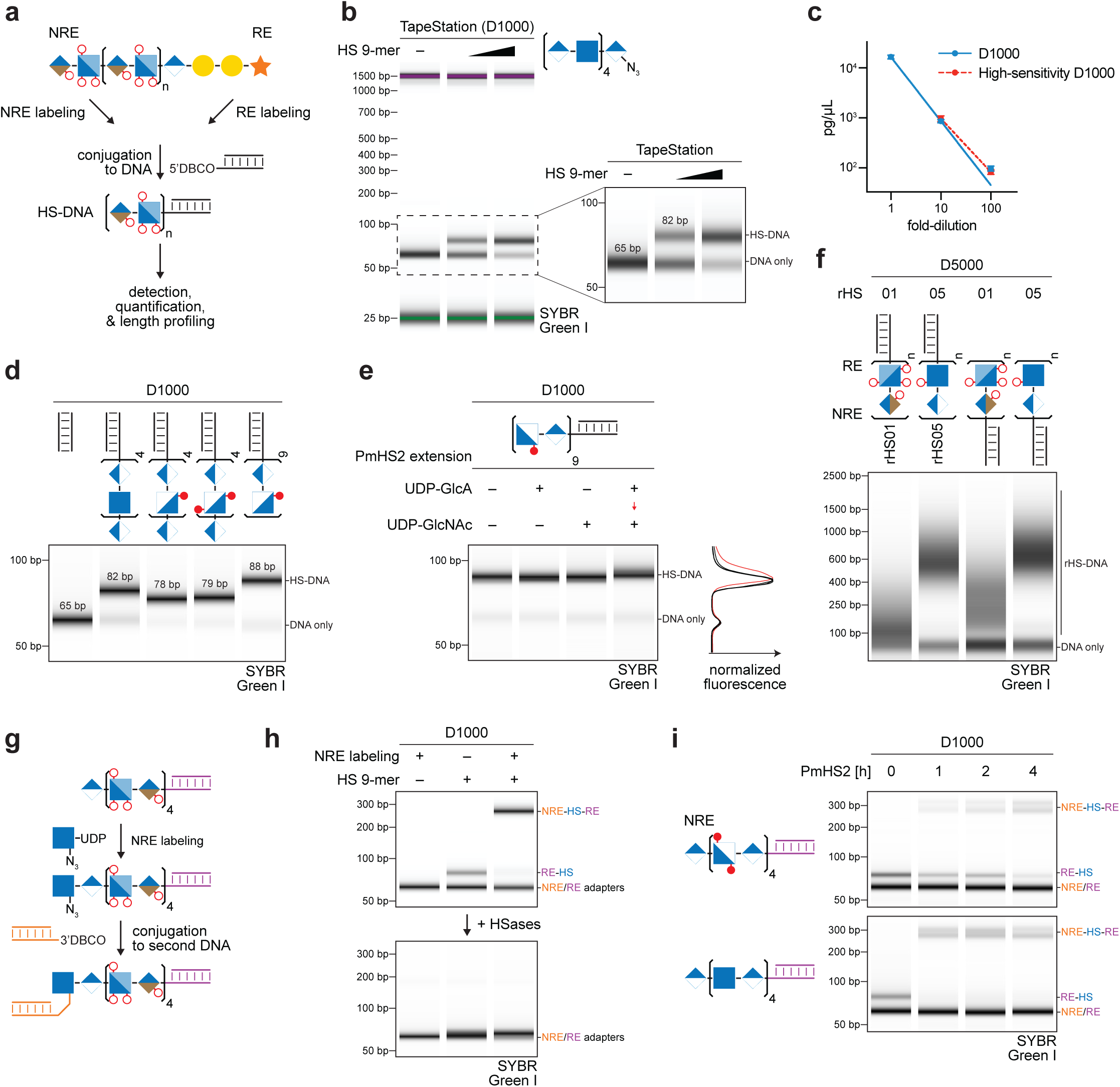
DNA conjugation reframes intact heparan sulfate analysis as a nucleic acid measurement. a. Schematic of HS-DNA conjugation. Azide-labeled HS chains are conjugated to DBCO-functionalized DNA by copper-free click chemistry (SPAAC), enabling detection, quantification, and length profiling on standard nucleic acid analysis platforms. b. TapeStation (D1000) analysis of a synthetic HS 9-mer conjugated to DNA. c. Quantification of HS-DNA detection sensitivity across a dilution series using D1000 and High-Sensitivity D1000 TapeStation assays. Curve represents a nonlinear regression fit to a log-log line model, with both axes displayed on logarithmic scales (GraphPad Prism). n = 3; bars indicate mean ± s.d. d. TapeStation (D1000) analysis of DNA conjugates of synthetic HS 9-mers of increasing sulfation and an 18-mer. e. TapeStation (D1000) analysis of PmHS2-mediated extension of an HS 18-mer-DNA conjugate with sequential monosaccharide additions. Right, normalized electropherograms. f. TapeStation (D5000) analysis of cell-derived rHS01 and rHS05 conjugated to a DNA adapter at either the RE or NRE. g. Schematic of DNA-HS-DNA chimera cloning. An RE-conjugated HS-DNA is enzymatically labeled at the NRE and conjugated to a second DNA adapter. h. TapeStation (D1000) analysis of chimera assembly with (+) or without (-) HS 9-mer and NRE labeling. Bottom, HSases digestion of each sample. i. TapeStation (D1000) time-course (0, 1, 2, 4 h) of PmHS2-mediated extension during chimera assembly on two synthetic HS 9-mers differing in sulfation near the NRE.

The apparent nucleic acid length of HS-DNA conjugates was determined by both chain length and sulfation. A synthetic 18-mer migrated slower than a 9-mer of comparable sulfation, while increasing sulfation across 9-mers reduced their apparent size, consistent with charge-dependent compaction observed in traditional HS gel electrophoresis^36^ (Fig. 2d). This length sensitivity was sufficient to resolve the addition of GlcA-GlcNAc disaccharide to a synthetic HS 18-mer by PmHS2, whereas addition of either monosaccharide alone produced no detectable shift (Fig. 2e). Native cell-derived HS conjugated to DNA produced broad migration profiles reflecting the polydisperse distribution of chain lengths, with overall similar, but distinct profiles depending on whether the DNA was conjugated at the RE or NRE (Fig. 2f and Figure S4b).

We extended the HS-DNA conjugate (DNA on the RE) into a DNA-HS-DNA chimera by NRE labeling and conjugation to a second DNA adapter (Fig. 2g). The chimera migrated like a ∼300 bp nucleic acid fragment despite containing only ∼120 bp of DNA, suggesting that the connecting HS segment significantly alters the migration through the capillary electrophoresis matrix (Fig. 2h). Chimera formation required both HS and PmHS2-mediated NRE labeling, and HSases digestion collapsed the chimera to its constituent DNA adapter. Cloning of fourteen synthetic 9-mers of different sulfation revealed that the standards with heavily sulfated residues near the NRE produced lower chimera yields (Figure S4c). Extension of the duration of PmHS2 labeling improved the yield of dual-end-labeled substrates, consistent with observed reduced PmHS2 activity toward highly sulfated NREs (Fig. 2i).

### Enzymatic translocation of DNA-HS-DNA conjugates through a biological nanopore

Given the linear, polyanionic, and DNA-terminal nature of our DNA-HS-DNA chimeras, we hypothesized they would be competent for translocation through DNA motor-based nanopores. Previous analysis of GAGs with solid-state and biological nanopores has been limited by the rapid, uncontrolled translocation of the polysaccharide chains^23^ (Figure S5a). We reasoned that flanking the HS with DNA adapters would thread the chimera more consistently through a biological nanopore with a DNA helicase motor (Fig. 3a).

**Figure 3.**
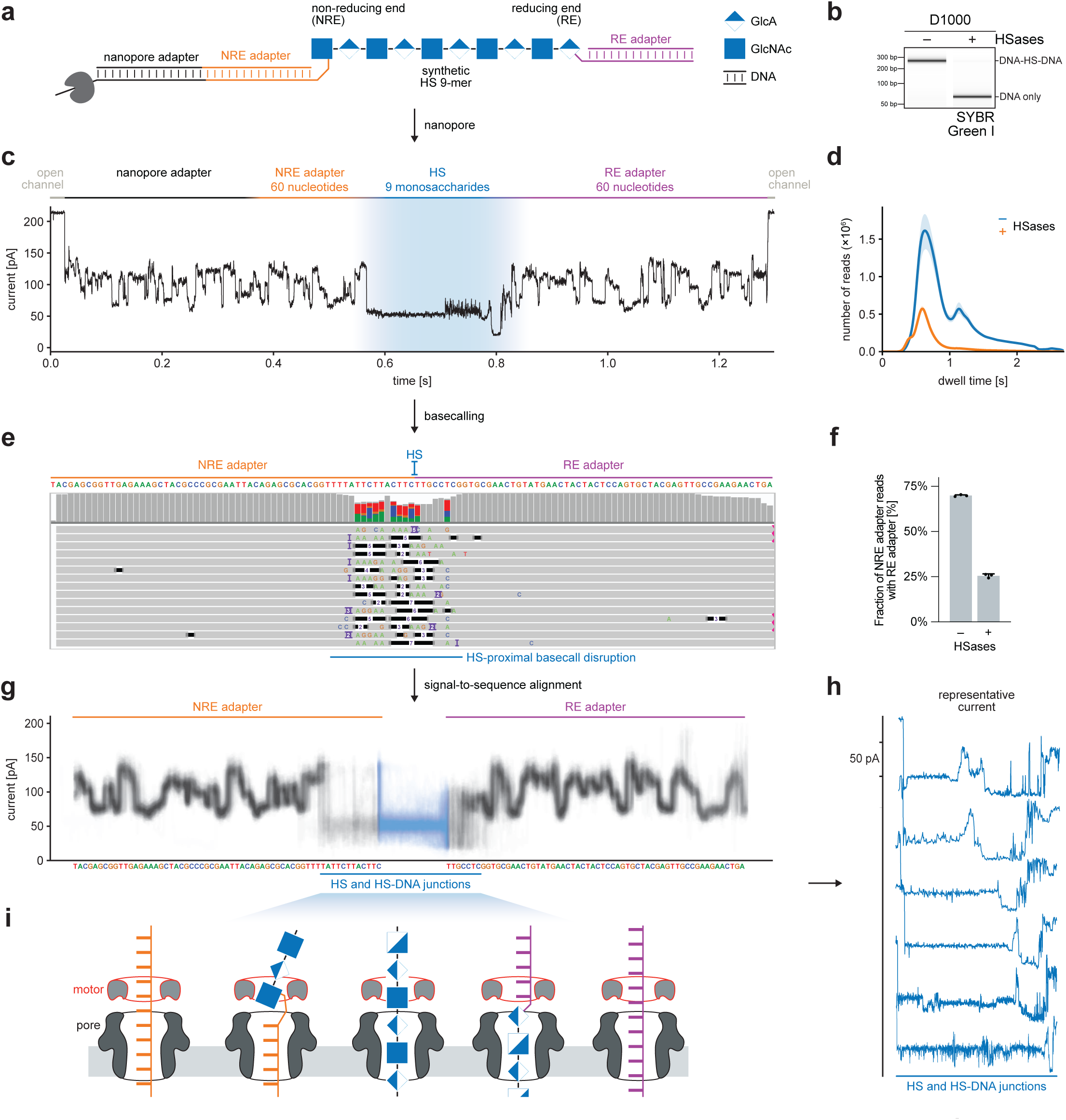
DNA-HS-DNA conjugates translocate through a motor-driven nanopore. **a.** Schematic of nanopore analysis of a synthetic DNA-HS-DNA chimera. A synthetic HS 9-mer flanked by 60-bp DNA adapters at the non-reducing end (NRE) and reducing end (RE) is ligated to a nanopore sequencing adapter bearing a motor protein. **b.** TapeStation (D1000) analysis of the DNA-HS-DNA chimera with (+) or without (-) HSases. SYBR electropherograms are uniformly normalized across the experiment. **c.** Representative raw current trace from a single DNA-HS-DNA translocation event. Sequential current regimes correspond to the open channel, nanopore adapter, NRE DNA adapter (60 nt, orange), HS segment (9 monosaccharides, blue), RE DNA adapter (60 nt, purple), and return to open channel. **d.** Strand-duration distributions for intact (blue) and HSases-digested (red) DNA-HS-DNA libraries. n = 3 replicates. **e.** IGV alignment of basecalled reads to an adapter reference containing the NRE and RE adapters. **f.** Fraction of NRE adapter-aligned reads that also contain an RE adapter alignment, with (+) or without (-) HSases digestion. n = 3; bars indicate mean ± s.d. **g.** Signal-to-sequence segmentation of dual-mapping reads. Raw current traces from 500 dual-mapping reads partitioned into NRE adapter, HS, and RE adapter regions using basecaller move-table information. **h.** Individual current traces spanning the HS and HS-DNA junction regions. Traces are normalized to a common length and vertically offset for visualization. The displayed region includes 12 nt of the NRE-proximal DNA adapter, the HS segment, and 7 nt of the RE-proximal DNA adapter. Scale bar, 50 pA. **i.** Model of DNA-HS-DNA chimera translocation through the motor protein and nanopore constriction.

We used the nanopore sequencing platform from Oxford Nanopore Technologies (ONT) and ligated the NRE adapter to an ONT DNA sequencing adapter bearing a helicase motor (Fig. 3a, S5b). We chose NRE-to-RE directionality so that we limit translocation initiation to only HS chains with PmHS2-labeled NREs. Complete translocation of the chimeric molecule would result in RE adapter alignment as an anchor for stringent selection of full-length reads. As a negative control, we treated DNA-HS-DNA chimeras with HSases to degrade the HS linkage between the two adapters. TapeStation analysis confirmed nearly quantitative collapse of the chimera band (Fig. 3b and Figure S5c).

Next, we adapted intact and HSases-treated DNA-HS-DNA chimeras using ONT DNA sequencing chemistry. We then aligned the resulting reads and achieved an average of 200,000 to 900,000 individual, aligned reads per flow cell for intact and HSases-treated conditions (**Table S1**). Ionic current traces from the intact DNA-HS-DNA library contained three visually distinct regions (Fig. 3c): i) an ionic current segment corresponding to the NRE adapter, ii) an ionic current segment corresponding to translocation of the HS chain; and iii) an ionic current segment corresponding to the RE adapter. We noted distinct ionic current amplitudes for the HS-associated segment compared to the flanking DNA regions, consistent with the expected architecture of the chimera. Strand duration distributions revealed a prominent population of long reads (0.5 - 2 s) in the intact library that was strongly depleted in the HSases-treated control (Fig. 3d).

To confirm that these ionic current traces correspond to the DNA-HS-DNA chimeras, the reads were basecalled using ONT Dorado software and aligned to a reference sequence using minimap2. The reference included concatenated NRE and RE DNA adapter sequences (Fig. 3e). The basecalling accuracy along dual-mapping reads decreased in the region proximal to the HS segment, consistent with HS-derived current patterns disrupting the basecaller’s sequence prediction within the HS context window. To assess the specificity of this alignment strategy for selecting dual-mapping reads, we calculated the fraction of reads that aligned to both the NRE and RE adapters (Fig. 3f). This analysis identified 70% of all reads as dual-labeled. The dual-mapping fraction decreased in HSases-digested samples to 26%, in which cleavage of the HS oligosaccharide is expected to separate the two DNA adapters, confirming that the dual-adapter reads originated from intact, continuously translocated DNA-HS-DNA chimeric strands.

To visualize HS-associated ionic current patterns across reads, we segmented the dual-mapping reads into NRE adapter and RE adapter regions using the emit-moves feature of the Dorado basecaller^37^ and aligned them to the reference NRE-RE sequence, leaving the middle of the current traces as putative HS region (Fig. 3g). Consistent with the IGV alignments, current traces showed robust alignment across DNA regions distal from the HS segment but increased heterogeneity near the HS. This disruption extended asymmetrically into the flanking DNA: approximately 12 nt on the NRE side and 7 nt on the RE side, defining a region we term the HS and HS-DNA junction window. Inspection of individual current traces within this window (Fig. 3h) revealed reproducible current-level transitions, but with variable step sizes relative to the more uniform steps observed in the flanking DNA regions (Fig. 3i, Figure S5d). This could be explained, in part, by the various interaction modes of the DNA-HS-DNA chimera molecules with the enzyme-nanopore complex as the helicase transitions from NRE DNA adapter to the HS chain and then to the RE DNA adapter. Previous work has documented a similar phenomenon - changes in helicase enzyme kinetics due to RNA modifications can be observed as distal ionic current changes that occur for ∼11 nucleotides 3′ of the strand in the nanopore sensor^38^.

### Nanopore ionic current signatures associated with synthetic heparan sulfate oligosaccharides

Having established that DNA-HS-DNA chimeras translocate through a motor-driven nanopore, we next assessed if the HS-associated ionic current features encode structural information related to sulfation and uronic acid composition. To test this, we synthesized a panel of 16 synthetic HS standards comprising fourteen 9-mers and two 18-mers with diverse sulfation modifications (Fig. 4a). Each standard was conjugated to DNA adapters (Figure S6a) and the resulting DNA-HS-DNA chimera was purified by UPLC (Figure S7a), barcoded with ONT adapters, pooled, and sequenced on a single nanopore flow cell.

**Figure 4.**
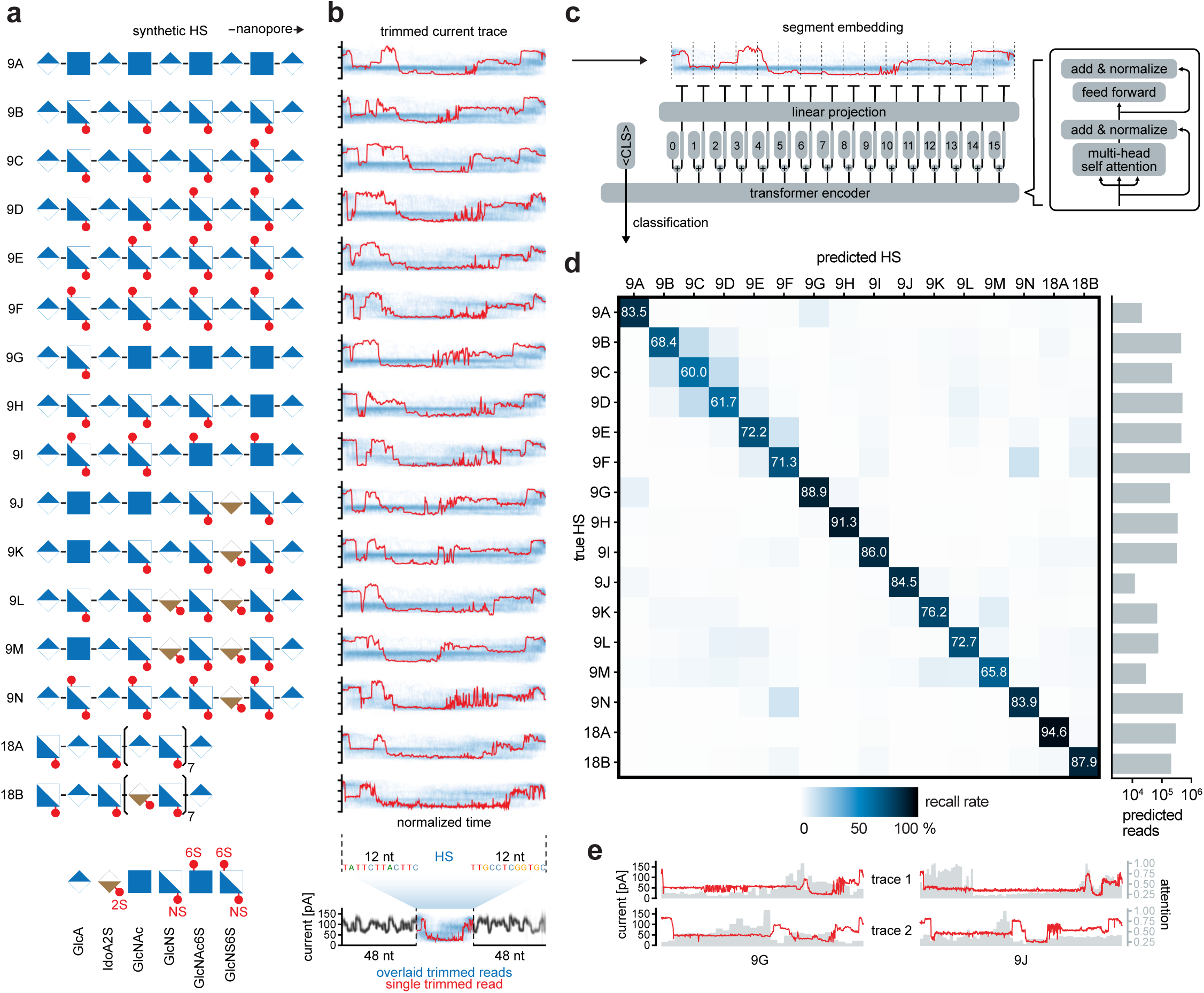
Nanopore current signatures fingerprint synthetic heparan sulfate oligosaccharides. a. Panel of synthetic HS standards for nanopore current fingerprinting. SNFG^45,46^ structures are shown for 16 standards, comprising fourteen 9-mers and two 18-mers, with sulfation positions annotated. Bottom, SNFG legend for HS monosaccharide building blocks with sulfation position annotations (2S, NS, 6S). b. Overlaid nanopore current traces for each HS standard. HS-associated current segments were trimmed with 12 nt of flanking DNA adapter sequence on each side. Blue, overlay of 100 reads per standard; red, representative single reads. Bottom, full-length trace for 18B indicating the trimmed region used for comparison. c. Architecture of the transformer encoder used for HS classification from raw nanopore current. Segmented current traces are positionally encoded, prepended with a classification (CLS) token, and passed through multi-head self-attention and feed-forward layers. d. Confusion matrix showing per-class recall (%) across the 16-standard HS panel. Right, total predicted reads per class (log scale). e. Representative trimmed current regions for HS 9G and 9J, with classifier attention weights mapped onto the current signal.

All 16 chimeras produced comparable read counts after demultiplexing, consistent with their equimolar content in the input (Figure S8a,b). Using the same basecalling and alignment strategy (Figure 3e), we identified dual-mapping reads for each standard (Figure S8b). Strand durations among the dual-mapped reads were comparable across standards (Figure S8c). To isolate the HS-derived signal, we trimmed each current trace to 12 nucleotides of flanking DNA adapter sequence on each side (Fig. 4b and Figure S8d). Overlaid current traces from 100 reads per standard (blue) revealed visually distinct current profiles, whereas individual reads (red) were more heterogeneous, consistent with the stochastic variation during motor-driven translocation^39^.

To classify HS chains from single-read current traces, we trained a transformer model on the trimmed ionic current segments (Fig. 4c), adapting an architecture previously used for nanopore-based tRNA classification^40^. Using 75% of the reads from the lowest abundance sample, the classifier achieved per-class recall ranging from 60.0% (9C) to 94.6% (18A) on a held-out test set. Thus, single-molecule traces derived from HS oligosaccharides produce discriminable current signatures despite read-to-read variability (Fig. 4d). Examining the “attention” of the model (i.e. the region within the ionic current with the highest model confidence for classification), we identified regions within the trimmed current that contributed most strongly to classification (Fig. 4e).

Further exploration of the confusion matrix (a tabular representation of machine learning-based classification) revealed specific off-diagonal misclassification patterns which were themselves informative, revealing chemical features the model could be distinguishing. Recall rates above 1% grouped structurally related standards (Figure S9). Several pairs showed symmetrical misclassification - 9A/9G and 9F/9N differ by a single sulfation group and were misclassified with one another. Within the 9K-9L-9M cluster, pairs differing by one monosaccharide (9K/9M, 9L/9M) showed elevated misclassification while the two-monosaccharide pair (9K/9L) did not, suggesting that the practical resolution of the present model could be at a single-monosaccharide difference. The 9B-9F cluster, which represents progressive 6-*O*-sulfation along a GlcNS backbone, had the highest misclassification rates, suggesting that 6-*O*-sulfation produces subtle changes in current fingerprints. Cross-cluster confusion between 9L-9M and 9B-9E further suggests that IdoA2S may generate ionic current features resembling those produced by an adjacent 6S group, likely owing to the conserved charge of the disaccharide bases. Together, these error patterns indicate that the transformer model learns chemically meaningful features and misclassifications track structural similarity rather than random class assignment.

### Native HS chains yield sulfation-dependent nanopore signatures

We next sought to extend this analysis to cell-derived heparan sulfate chains, which are longer and more heterogeneous than our synthetic standards, but compositionally defined by disaccharide abundance determined by heparin lyase treatment. Having established methods to selectively mono-label native HS chains at each end with a DNA adapter, to visualize the HS-DNA conjugates, and to demonstrate translocation of HS-DNA chimeras through a nanopore, we combined these steps to examine the ionic current signatures of intact heparan sulfate chains using the same end-adaptation strategy (Fig. 5a). We selected two cell-derived HS chains that differ in both sulfation and chain length as model polymers (Fig. 5b). rHS01, derived from wild-type CHO cells, represents a chain with a native sulfation pattern and motifs, whereas rHS05 is markedly undersulfated, retaining only minimal 6-*O*-sulfation.

**Figure 5.**
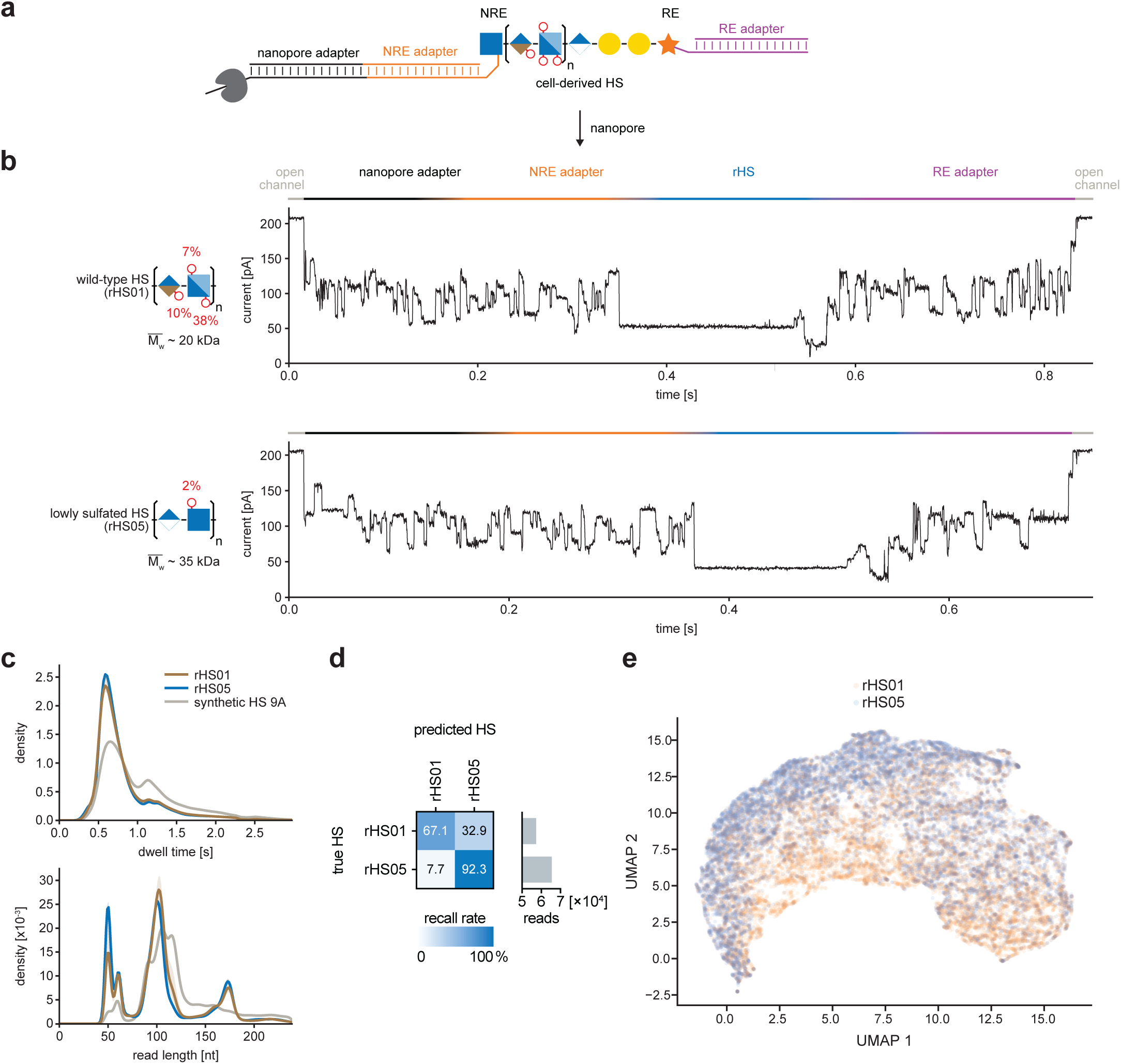
Native heparan sulfate chains produce sulfation-dependent nanopore signatures. **a.** Schematic of nanopore analysis of a cell-derived DNA-rHS-DNA chimera. Cell-derived rHS flanked by 60-bp DNA adapters at the non-reducing end (NRE) and reducing end (RE) is ligated to a nanopore sequencing adapter bearing a motor protein. **b.** Representative raw current traces (one per class) from single DNA-rHS-DNA translocation events for CHO HS (rHS01) and lowly sulfated (rHS05) heparan sulfate chains. Sequential current regimes correspond to the open channel, nanopore adapter, NRE DNA adapter (60 nt, orange), HS segment (blue), RE DNA adapter (60 nt, purple), and return to open channel. **c.** Strand-duration distributions for intact DNA-rHS-DNA libraries compared with the strand duration of an unsulfated synthetic HS 9-mer. n = 3 replicates. **d.** Confusion matrix showing per-class recall (%) for binary classification of rHS01 and rHS05 nanopore current traces. Right, total predicted reads per class (log scale). **e.** UMAP projection of per-read signal features extracted from trimmed current segments of rHS01 and rHS05 dual-mapping reads. Features include median current, standard deviation, kurtosis, skewness, and dwell time. Each point represents a single translocation event, colored by class.

We first functionalized the reducing ends by hydroxylamination and conjugated them to a DNA adapter (Figure S10a). Because native HS and DNA co-purify under the silica-based purification method used here - and likely under alternatives such as DEAE-based anion exchange - the amount of input HS had to be controlled at the reaction step rather than removed downstream. As little as 20 pmol of rHS was sufficient to convert the majority of the DNA adapter despite the second-order kinetics of the copper-free (SPAAC) click reaction. Residual unreacted HS was then capped with CF647-DBCO and excess DNA adapter with sodium azide, blocking the reactive group on each species. The resulting HS-DNA conjugate was modified at the non-reducing end as described above and conjugated to a second DNA adapter; these capping steps enforce the correct NRE-to-RE directionality of the final reads. Finally, the DNA-HS-DNA conjugates - together with low-abundance RE-conjugated and unconjugated HS species – were UPLC-separated from free DNA adapters to maximize the fraction of informative reads on the pore (Figure S10b, c).

We then sequenced the DNA-modified rHS chains on the nanopore. Individual reads resolved into the expected succession of segments - open channel, nanopore adapter, NRE adapter, the HS chain, and the RE adapter - with the HS region appearing as a sustained low-current clearly distinct from the flanking DNA adapters (Fig. 5b). The raw traces themselves resembled those obtained with synthetic standards. While the rHS chains had nearly identical dwell time and the apparent read length after alignment, there were minor but specific differences in these distributions compared to the synthetic standards (Fig. 5c). Next, we focused the analysis on trimmed HS-proximal current regions which yielded class separation: rHS05 reads were recovered with high recall (92.3%), whereas rHS01 reads were recovered at 67.1%, with roughly a third classified as rHS05 (Fig. 5d). This asymmetry is consistent with the underlying composition: undersulfated rHS05 is comparatively uniform and occupies a tighter region of current space, whereas wild-type rHS01 is structurally heterogeneous with extended regions lacking any modifications (NA domains). An examination of ionic current features using dimensionality reduction (in UMAP space) revealed clusters that had both regions of demixing and regions with overlap between the two classes (Fig. 5e), consistent with the heterogeneity of both substrates in sulfation and length. Together, we measured > 5,700,000 individual molecules of native heparan sulfate from mammalian cells.

## Discussion

In this work, we developed strategies to purify and analyze intact heparan sulfate chains which enabled coupling of DNA to the chains for analysis by DNA-sensing technologies. Glycosaminoglycans, including HS and other glycans, are typically analyzed by bottom-up methods in which the chains are cleaved to disaccharides by enzymatic or chemical means and the disaccharides are analyzed by UV absorption, fluorophore tagging or mass spectrometry^7,27^. Their heterogeneity and length and the lability of sulfate esters when subjected to ionization generally prevent their sequencing in full-length form. GAGs are linear, sulfated polyanions, properties shared with nucleic acids and exploited here to enable their analysis with nucleic acid tools. By conjugating HS to DNA, we convert it into a substrate compatible with the nucleic acid tools, enabling high-sensitivity size characterization by gel electrophoresis at inputs far below the detection limits of conventional glycan staining (e.g. with Alcian Blue). More broadly, DNA conjugation transforms the manipulation of HS chains into a process analogous to molecular cloning; a DNA-HS(-DNA) chimera can be purified, resolved, and sequenced using infrastructure built for nucleic acids. Together our work provides a deployable framework to manipulate and sense HS chains, as well as other GAGs, across a wide variety of systems and settings.

The biological context in which HS operates is long (100+) monosaccharides in series tethered to a proteoglycan core. Conventional HS analysis derives its specificity from degradation - heparin lyases selectively cleave HS into disaccharides, and the resulting fragments are identified by their enzymatic origin^31^. This leaves other “contaminating” GAG chains (and co-purifying molecules) intact and thus not analyzed. However, establishing a strategy to profile intact HS chains requires an alternative basis for specificity that does not depend on degradation. Our dual-labeling strategy addresses this by combining chemoenzymatic NRE labeling, which selects for HS-like substrates of PmHS2, with chemical RE labeling via hydroxylamination. While PmHS2 could be expressed at higher levels and was more active than human EXT1/2, we still observed lower-efficiency labeling of the NRE as compared to the RE, suggesting potential specificity to particular sulfation patterns at or near to the NRE. New enzyme design or engineering efforts could facilitate application of a wider array of HS substrates into our chemoenzymatic framework.

Single-molecule sensing provides unique views into the molecular features of the analytes detected and given the importance of GAGs, this has been of interest for some time. Previous efforts to detect GAGs using nanopore sensing leveraged both solid-state^20^ and biological pores^19,21^, applying unmodified GAG chains to the pores. These approaches yielded uncontrolled translocation and generally relied on sample purity to interpret current traces as being relevant to the measurement of interest. By installing two defined DNA adapters onto the HS chains, we resolve, at least in part, the translocation control issue by leveraging motor-protein-controlled threading through the biological pore. This approach was conceptually paralleled by recent work using DNA-peptide conjugates to engage DNA motors^41^. Critically, because we use enzymes that only modify HS, we can use computational selection of current traces bearing both DNA adapter signatures to ensure full-length, orientation-resolved translocation that isolates the current region corresponding only to the HS chain for downstream classification.

Beyond demonstrating that complete translocation was possible for a wide diversity of HS chains, we documented that individual and precisely sulfated HS chains can be directly classified at the single-molecule level using only nanopore ionic current data. Our transformer model-based classifier achieved robust discrimination across 16 synthetic standards which varied across the number and position of sulfates, uronic acid composition, and length. What was particularly revealing from this analysis is where the transformer misclassified individual HS chains: the structure of its errors is chemically informative. Misclassifications were concentrated among structurally similar oligosaccharides, consistent with ionic current signatures that reflect the underlying sulfation pattern rather than nonspecific variation. Expanding the synthetic standard library to encompass additional sulfation permutations and chain lengths will be essential for decoding current signals from longer, native HS chains - a setting where no orthogonal method for full-length sequence determination currently exists. Importantly, because DNA-based technologies enable barcoded multiplexing, scaling up training sets and expanding into many biological samples in parallel can be implemented easily within the processing and analysis framework.

That the glycopolymers in the ECM, especially GAGs, encode regulatory logic based on precise compositions and sequences is long established. Yet GAG structural analysis generally remains limited to disaccharide-level measurements, lagging decades behind nucleic acid and protein sequencing. The disaccharide composition of a GAG chain provides limited insight into structure-function relationship and at best provides a measurement comparable to ACGT content of a DNA strand. Our nanopore-based measurements demonstrate discriminatory power on synthetic HS chains. Extending this resolution to native chains would begin to close the decades-long gap between GAG analysis and nucleic acid sequencing.

## Methods

### Heparan sulfate sources

Synthetic heparan sulfate standards (**Table S2**, Glycan Therapeutics) were resuspended to 500 µM in UltraPure water (10977015, Thermo Fisher Scientific) based on the manufacturer’s reported mass and molecular weight. Commercial cell-derived heparan sulfate (**Table S2**, TEGA Therapeutics) was resuspended to 200 µM in UltraPure water based on the manufacturer’s reported mass and average molecular weight (cell-derived heparan sulfate chains are a distribution of sizes).

### Heparan sulfate purification strategy

Heparan sulfate chains and HS-nucleic acid conjugates were purified on silica spin columns (Zymo-Spin I, C1003, or Zymo-Spin III, C1005; Zymo Research). The silica columns were pre-conditioned by adding 400 µL UltraPure water followed by centrifugation at 10,000 x g for 40 sec. HS-containing samples were mixed with 2 volume equivalents of RNA binding buffer (R1013-2, Zymo Research) and briefly vortexed. Subsequently, 6 volume equivalents of 100% ethanol (4068, Greenfield Global) were added, and the samples were vortexed thoroughly.

The resulting solution was applied to the pre-conditioned Zymo column and centrifuged at 10,000 x g for 20 sec, and the flow through was discarded. The column was washed twice with 650 µL of 80% ethanol (prepared by dilution of 100% ethanol with UltraPure water; 4068, Greenfield Global), centrifuging the first wash for 20 seconds and the second wash for 40 seconds to ensure complete removal of residual liquid. The bound material was eluted by adding UltraPure water to the column, incubating for 5 min at room temperature, and centrifuging at 10,000 x g for 40 sec. The elution step was repeated once.

Elution volumes were adjusted depending on downstream applications. For Zymo-Spin I columns, samples were eluted twice with 6.7 µL UltraPure water, yielding a combined eluate of ∼11 µL after accounting for column retention losses; this volume was used for small-scale biochemical reactions. When subsequent lyophilization was performed, the bound material was eluted twice with ≥10 µL UltraPure water.

### Silica column binding capacity assay using radiolabeled heparin

End-labeled tritiated heparin was prepared by sodium borotritide reduction of unfractionated therapeutic heparin (950611, McKesson), adapting a previously described method for heparin oligosaccharides^42^. [^3^H]heparin (2,000 cpm) was mixed with increasing amounts of unlabeled heparin (0-100 µg) and purified using a Zymo I column as described above. Following elution, each sample was transferred to a vial containing 5 mL Ultima Gold™ (6013321, Revvity). Radioactivity was measured using a HIDEX 300SL liquid scintillation counter, with counts acquired for 5 min.

### Heparinases digestion

Digestion of heparan sulfate was performed with heparin lyase I (HSase I; P0735, NEB), heparin lyase II (HSase II; P0736, NEB), and heparin lyase III (HSase III; P0737, NEB). Heparan sulfate was combined in a PCR tube to a final volume of 10 µL containing 12 U HSase I (1 µL stock), 4 U HSase II (1 µL stock), 0.7 U HSase III (1 µL stock) in 1x Heparinase Reaction Buffer (20 mM Tris-HCl pH 7, 100 mM NaCl, 1.5 mM CaCl_2_; 1 µL of 10x buffer; B0735, NEB). The mixture was incubated for 1 h at 30°C in a thermocycler. Negative control reactions lacking enzymes were incubated in 1x Heparinase Reaction Buffer. For direct in-gel analysis, the corresponding gel loading buffer (see “*Gel analysis of heparan sulfate*”) was added to each sample, and the mixture directly loaded onto the gel. Otherwise, the reaction mixture was purified using a Zymo I column as described above.

### Reducing end-labeling of heparan sulfate

The reducing end of heparan sulfate chains was labeled by hydroxylamination using a previously reported procedure^35^. Specifically, heparan sulfate chains (200 pmol unless otherwise noted) were combined in a PCR tube to a final volume of 20 µL containing 10 mM aminooxy-PEG_3_-azide (BP-22704, BroadPharm), 100 mM aniline (242284, Sigma) as catalyst, and 500 mM sodium acetate (pH 4.5, S0292, Teknova). The reaction mixture was incubated for 1 hour at 37°C in a thermocycler. Upon completion, labeled HS chains were purified using a Zymo I column as described above.

### Non-reducing end labeling of heparan sulfate

The non-reducing ends (NRE) of heparan sulfate chains were chemoenzymatically labeled with an azide handle using either the EXT1/EXT2 heterodimer (EXT1/2; 8567-GT-020, Bio-Techne) or the *Pasteurella multocida*-derived Heparosan Synthase 2 (PmHS2). Unless otherwise noted, native heparan sulfate chains or synthetic 18-mer HS oligosaccharides were first subjected to NRE repair by extension with UDP-GlcA, followed by NRE labeling using UDP-GlcNAz to install the azide functionality.

For PmHS2-mediated labeling, heparan sulfate was combined in a final volume of 20 µL containing 40 mM Tris-HCl (pH 7.5, T1075, Teknova), 4 mM MnCl_2_, 4 mM MgCl_2_, 100 µM UDP-GlcNAz (HY-145934A, MedChemExpress) or 100 µM UDP-GlcA (U6751, Sigma), and 450 µg/mL of PmHS2, using a modified version of previously reported reaction conditions^43^. The reaction was incubated at 32°C on a thermocycler for 1 h. The reaction mixture was purified using a Zymo I column as described above.

For EXT1/2-mediated labeling, heparan sulfate was mixed in a final volume of 20 µL containing 25 mM Tris-HCl (pH 7.5, T1075, Teknova), 150 mM NaCl, 10 mM MnCl_2_, 100 µM UDP-GlcNAz (HY-145934A, MedChemExpress), and 1 µL of EXT1/2 (0.3 mg/mL; dependent on manufacturer’s lot concentration), using a modified version of previously reported reaction conditions^26^. The reaction was incubated at 37°C on a thermocycler for 12 h. The reaction mixture was purified using a Zymo I column as described above.

### Copper-free click labeling of heparan sulfate

For in-gel detection, azide-modified HS labeled at either the reducing end (RE) or non-reducing end (NRE) was subjected to copper-free click (SPAAC) with IRDye 800CW-DBCO. Purified azide-modified HS eluate (∼11 µL from a Zymo-Spin I column) was mixed with 1 µL of 10× PBS (25-507X, Genesee Scientific) and 1 µL of 1 mM IRDye 800CW-DBCO (929-55000, LI-COR Biosciences). The reaction was incubated at 37°C for 30 min, followed by purification using a silica spin column as described above.

### Gel analysis of heparan sulfate

Heparan sulfate, including nucleic acid-HS conjugates, was resolved by either native polyacrylamide gel electrophoresis (native PAGE) or AnyKD SDS-PAGE gels (5671124, Bio-Rad) and detected using either Alcian Blue (J60122.09, Thermo Fisher Scientific) staining or SYBR Gold (S11494, Thermo Fisher Scientific) for nucleic acid-containing species.

Unless otherwise noted, samples were resolved on 10% native PAGE gels. Gels were prepared by mixing 6.5 mL UltraPure water, 1 mL 10x UltraPure TBE Buffer (1 M Tris, 0.9 M boric acid, and 0.01 M EDTA; 15581044, Thermo Fisher Scientific), and 2.5 mL 40% acrylamide/bisacrylamide solution (1610146, Bio- Rad). Polymerization was initiated by sequential addition of 80 µL of 10% (w/v) ammonium persulfate solution (A3678, Sigma) and 4 µL UltraPure TEMED (15524010, Thermo Fisher Scientific). The solution was poured into an Invitrogen casting station and allowed to polymerize for at least 20 min. Prior to sample loading, gel was pre-run at 5 W for 5 min in 1x UltraPure TBE buffer. Samples were mixed with BlueJuice loading buffer (10816015, Thermo Fisher Scientific) to a final concentration of 1x, loaded into the wells, and electrophoresed at 7 W for 30 min. In the absence of a commercially available heparan sulfate size ladder, a 25 bp DNA step ladder (G4511, Promega) was used as a migration reference under native PAGE conditions.

Following electrophoresis, the gel was briefly rinsed with Milli-Q water and stained with 1x SYBR Gold in 1x PBS for 10 min at room temperature. Gels were then rinsed with Milli-Q water and imaged on a Bio-Rad imaging system. Unlabeled HS and other acidic polysaccharides were detected using Alcian Blue staining as reported previously^44^. Briefly, a 0.1% (w/v) Alcian Blue solution in 2% acetic acid was prepared by dissolving 30 mg Alcian Blue dye in 29.4 mL UltraPure water supplemented with 600 µL of glacial acetic acid (A6283, Sigma). Gels were stained for 1 h, destained in 2% acetic acid for 15 min, and further destained in Milli-Q water overnight. Alcian Blue-stained gels were imaged on a Bio-Rad imaging system using the Coomassie Blue detection channel and band intensities were quantified from the raw images using Image Studio software (LI-COR Biosciences).

For detection of IR800-labeled heparan sulfate on AnyKD SDS-PAGE gels, samples were mixed 1:1 with formamide, loaded directly into the wells, and electrophoresed at 150 V for 30 min in 1x Tris-glycine-SDS running buffer (BP-150, Boston BioProducts). A Chameleon Duo pre-stained ladder (928-60000, LI-COR Biosciences) was included as a migration reference. Following electrophoresis, gels were briefly rinsed with Milli-Q water and imaged on a LI-COR Odyssey infrared imaging system. For in-gel protein visualization (e.g. heparinases), the gels were stained with Coomassie Blue (AS001000, Bulldog Bio) for 1 h at room temperature and destained in Milli-Q water overnight. Coomassie Blue-stained gels were imaged on a Bio-Rad imaging system.

### TapeStation analysis

DNA and DNA-HS conjugates were analyzed using D1000, High Sensitivity D1000, or D5000 ScreenTape assays on a TapeStation system (Agilent Technologies), following the manufacturer’s recommended protocols. ScreenTape types were selected based on the expected size range and concentration of the analyzed constructs. As an example, for the D1000 assay, 1 µL of sample was mixed with 3 µL D1000 Sample Buffer in an optical 8-tube strip, vortexed at 2,000 rpm for 1 min, and briefly centrifuged. Samples were then loaded onto the TapeStation instrument for analysis. Unless otherwise noted, the resulting gel electropherograms are displayed with per-sample intensity scaling.

### DNA adapter annealing and UPLC purification

DNA duplex adapters bearing a dibenzocyclooctyne (DBCO) modification on either the 3’ or 5’ terminus were synthesized by IDT (**Table S3**). Forward strands containing the DBCO modification and complementary reverse strands lacking DBCO were prepared as 100 µM stocks and mixed at a 6:10 molar ratio. Duplex annealing was performed in a thermocycler using the following temperature program: 90 °C 30 sec, 70 °C 1 min, 60 °C 1 min, 50 °C 1 min, 37 °C 1 min, and 25 °C 1 min.

Following annealing, the duplex mixture was diluted with HPLC-grade acetonitrile to a final concentration of 75% (v/v) and purified by ultra-performance liquid chromatography (UPLC) on a GTxResolve Premier BEH Amide column (300 Å, 1.7 µm, 2.1 x 100 mm). Separation was performed using a linear gradient from 60% to 90% solvent A over 10 min at a flow rate of 0.5 mL/min (solvent A: 100 mM NH_4_OAc in water, 09689, Sigma; solvent B: acetonitrile, 34851, Sigma).

The eluted fraction corresponding to the annealed duplex was dried under vacuum in a SpeedVac for 1 h at room temperature and subsequently lyophilized overnight. The lyophilized material was further purified using a Zymo I column as described above, diluted to final concentration of 10 µM, aliquoted and stored at −80°C.

### Conjugation of synthetic standards to DNA adapters

HS-DNA and DNA-HS-DNA conjugates of azide-tagged synthetic HS oligosaccharides (Glycan Therapeutics, **Table S2**) were generated through stepwise chemical and enzymatic modification with DBCO-modified dsDNA adapters.

For synthetic 9-mers, copper-free click coupling (SPAAC) was performed by combining 1 µL of 500 µM azide-modified HS oligosaccharide with 1 µL of 10 µM 5’DBCO-modified dsDNA adapter and 1 µL of 10x PBS (25-507X, Genesee Scientific). For synthetic 18-mers, 5 µL of each 50 µM stock were combined with 5 µL of 10 µM 5’DBCO-modified dsDNA adapter and 5 µL of 10x PBS. The mixture was vortexed, briefly spun down, and incubated at 37°C for 30 min in a thermocycler. The residual DBCO-reactive dsDNA adapter was quenched by adding 1 µL of 10 mM sodium azide (NaN_3_; S2002, Sigma) followed by incubation at 37°C for 10 min. Reactions were brought to 20 µL with UltraPure water and purified using a Zymo-I column as described above. The eluate contained the desired DNA-HS conjugate together with any remaining unreactive or quenched DBCO-modified dsDNA adapter.

To generate DNA-HS-DNA, the NRE of the DNA-HS conjugate was chemoenzymatically labeled with an azide using PmHS2 and UDP-GlcNAz as described above. For even-numbered chains (e.g. synthetic 18-mers) and for native HS chains, NRE repair was performed prior to azide installation by extension with UDP-GlcA for 12 h, followed by PmHS2-mediated incorporation of UDP-GlcNAz for 4 h. After each enzymatic step, the reaction was purified using a Zymo-I column as described above. The NRE-azide-labeled DNA-HS conjugate was lyophilized and subjected to a second SPAAC reaction by combining the material with 1 µL of 10 µM 3’DBCO-modified dsDNA adapter and 1 µL of 10x PBS, followed by incubation at 37°C for 30 min. Reactions were brought to 20 µL with UltraPure water and purified using a Zymo-I column as described above to yield DNA-HS-DNA conjugates.

### Conjugation of cell-derived HS to DNA adapters

rHS-DNA and DNA-rHS-DNA conjugates were generated by stepwise chemical and enzymatic modification of cell-derived HS (TEGA Therapeutics) at the reducing end (RE) and non-reducing end (NRE) with DBCO-modified dsDNA adapters.

Samples (200 pmol) of rHS01 (4 µg) or rHS05 (7 µg) (molecular mass values for the chains provided by TEGA Therapeutics) were hydroxylaminated and purified on a Zymo column as described above. 1 µL of eluate (∼20 pmol) was conjugated to a 5’DBCO-modified dsDNA adapter via SPAAC by combining it with 1 µL of 10 µM adapter and 1 µL of 10× PBS, followed by incubation at 37 °C for 1 h. The reaction was purified on a Zymo column. Unreacted azide groups on excess rHS were quenched by the addition of 1 µL of 10 mM CF647-DBCO (96117, Biotium) and 1 µL of 10× PBS, incubation at 37 °C for 1 h, dilution to 80 µL, and Zymo column purification. Unreacted DBCO groups on excess adapter were quenched by the addition of 1 µL of 10 mM sodium azide (S2002, Sigma-Aldrich) and 1 µL of 10× PBS under the same conditions.

To generate DNA-rHS-DNA conjugates, the NRE of the purified rHS-DNA conjugate was chemoenzymatically labeled with an azide handle. The NRE was first repaired by extension with UDP-GlcA using PmHS2 for 12 h, followed by PmHS2-mediated incorporation of UDP-GlcNAz for 4 h. Each enzymatic step was followed by Zymo column purification. The NRE-azide-labeled rHS-DNA conjugate was lyophilized, then conjugated to a 3’DBCO-modified dsDNA adapter via SPAAC (1 µL of 10 µM adapter, 1 µL of 10× PBS, 37 °C, 1 h). The reaction was diluted to 80 µL and purified on a Zymo column. DNA-rHS-DNA conjugates were further purified from unconjugated adapters by UPLC as described below.

### UPLC separation of HS-DNA conjugates

HS-DNA and DNA-HS-DNA conjugates were mixed with HPLC-grade acetonitrile (34851, Sigma-Aldrich) to a final concentration of 75% (v/v) and purified by UPLC using the column and solvent system described above. DNA-HS-DNA conjugates of synthetic HS were separated starting with an isocratic hold at 60% solvent A for 3 min, which elutes the free dsDNA adapters and HS-DNA mono-conjugates, followed by a linear gradient from 60% to 80% solvent A over 5 min (Figure S7). DNA-rHS-DNA conjugates of cell-derived HS were separated starting 60% solvent A for 5 min which elutes the free dsDNA adapters, followed by a linear gradient from 60% to 100% solvent A over 1 min (**Figure S10**). Eluted fractions were collected, dried under vacuum in a SpeedVac for 1 h at room temperature, lyophilized overnight, and further purified using a Zymo-I column as described above.

### ONT adapter ligation and nanopore sequencing

HS-DNA and DNA-HS-DNA conjugates were first 5’ phosphorylated using T4 Polynucleotide Kinase (T4 PNK) to enable subsequent adapter ligation. Briefly, HS-DNA conjugates (10 pmol or less) were brought to a final volume of 30 µL containing 20 U T4 Polynucleotide Kinase (2 µL stock; M0201, NEB), 1 mM ATP (P0756, NEB), and 1x T4 PNK buffer (70 mM Tris-HCl, 10 mM MgCl_2_, 5 mM DTT, pH 7.6; B0201, NEB). The reactions were incubated at 37°C for 30 min and subsequently purified using a Zymo I column as described above.

5’ phosphorylated HS-DNA and DNA-HS-DNA conjugates were then processed using the Native Barcoding Kit 24 V14 from Oxford Nanopore Technologies according to the manufacturer’s protocol. Briefly, barcoding and sequencing adapters were ligated to the phosphorylated DNA termini using T4 DNA ligase, after which the ligation reactions were cleaned up following the kit recommendations. Prepared libraries were loaded onto nanopore flow cells and sequenced using standard PromethION run parameters.

### Basecalling, read alignment, and current trace isolation

Ionic current traces were basecalled using Dorado 1.0.0 (GitHub) and aligned to a reference containing the two DNA adapters.

### Chain Classification

To characterize the sulfation diversity of HS chains across samples, we developed a supervised classification framework to assign individual nanopore current traces to their corresponding HS chain class. Each trimmed current segment from a dual-mapping read served as input, and the model was trained to predict the identity of the HS standard from which the read originated.

### Preprocessing and Signal Trimming

Raw ionic current signals from dual-mapping reads were preprocessed to remove barcode-derived sequence contributions prior to classification. Because barcodes are used to distinguish samples within a multiplexed sequencing run but do not correspond to HS-derived current, their inclusion would introduce confounding features. To remove barcode signals, each read was aligned to the non-barcoded reference sequence, and basecaller move tables were used to identify the signal region corresponding to the reference. Signal segments outside this correspondence were discarded. The resulting traces were further trimmed to retain the HS-associated current segment flanked by 12 nucleotides of the NRE and RE DNA adapter sequence on each side, yielding a standardized input window free of barcode contamination that captures both the HS segment and the HS–DNA junction regions.

### HS Chain Classification

Trimmed current traces were used to train a transformer-based encoder for HS classification, adapting an architecture previously applied to nanopore-based tRNA identification^40^. To ensure balanced representation across classes, we identified the class with the fewest reads and used this count to define a uniform per-class sample size for training and validation. For each class, reads were randomly assigned to training (75%) and validation (5%) subsets based on this uniform count, with all remaining reads allocated to the test set. These subsets were fixed and saved to ensure reproducibility across experiments. The same balancing and splitting strategy was applied consistently across all classification tasks described below.

As preliminary validation steps, we first trained an orientation classifier to distinguish forward from reverse-complement reads, confirming that strand directionality is encoded in the current signal. We subsequently trained a binary classifier to discriminate synthetic HS-derived traces from native, cell-derived HS traces, verifying that the model captures biologically meaningful signal differences. The final 16-class model was evaluated on the held-out test set, with classification performance assessed by per-class recall and summarized as a confusion matrix across all 16 standards.

### Misclassification Analysis

To assess whether classification errors reflected underlying chemical similarity among HS standards, we examined off-diagonal entries in the confusion matrix. Standard pairs with symmetrical misclassification rates exceeding 1% recall were identified and compared with respect to their structural differences, including the number and position of sulfation modifications and monosaccharide composition. This analysis enabled grouping of structurally related standards based on their confusion profiles and provided insight into the chemical features that produce similar or overlapping current signatures.

### Interpretability via Attention Analysis

To identify which regions of the current trace contribute most to classification, we examined the self-attention weights of the trained transformer. Attention maps were aligned to the underlying signal and nucleotide positions using basecaller move tables, enabling mapping of high-attention regions back to specific positions along the trimmed trace. For each standard, positions with the highest attention weights were compared against the known HS and HS-DNA junction boundaries to determine whether the model preferentially attends to HS-derived current, adapter-proximal current, or both. This analysis identified the molecular features of the chimera most informative for distinguishing individual standards as they engage the motor protein and nanopore constriction.

## Data Availability

Raw nanopore trace data are available upon request. Any additional information is available from the corresponding contact upon request.

## Code Availability

This paper reports new code related to analysis of Nanopore-based HS translocation. The code is available via GitHub at https://github.com/genometechlab/HS-nano-seq.

## Acknowledgments

We thank the members of the Jain and Flynn laboratories for constructive feedback, Eric Schmidt, Kaori Oshima, and Lauren Pepi for useful discussions, Eliezer Calo for providing Alcian Blue reagent, Fernando Camargo and Xugeng Liu for providing heparin, and the Kean lab for the use of their TapeStation device. This work was supported by grants from the Rita Allen Foundation (R.A.F.), Yosemite (R.A.F.), the Scleroderma Research Foundation (R.A.F.), the G. Harold and Leila Y. Mathers Charitable Foundation (R.A.F.), Cancer Research Institute (CRI14279; R.A.F.), and R01GM145913 (to J.D.E.). R.A.F. is the Bakewell Foundation-Rachleff Innovator of the Damon Runyon Cancer Research Foundation (DRR-74-23).

## Author Contributions

P.H. and R.A.F. conceived the project. R.A.F. and M.J. supervised and obtained funding for the project. P.H. designed and performed the majority of the experiments. P.H. and S.K.R. designed and performed UPLC experiments. P.H. and T.T. designed and performed nanopore experiments. G.S. and J.L. synthesized synthetic HS standards and expressed PmHS2. P.O. and J.D.E. designed and performed radioactive heparin experiments. P.D.K. and M.J. developed computational methods and analyzed nanopore data. P.H., P.D.K., M.J., and R.A.F. wrote the manuscript. All authors discussed the results and revised the manuscript.

## Competing Interests

R.A.F. is a stockholder of ORNA Therapeutics. R.A.F. is a board of directors member and stockholder of Blue Planet Systems. R.A.F., P.H., and M.J. have filed a patent related to HS-DNA conjugation for analysis using DNA sensing platforms. M.J. is a consultant to ONT and has received reimbursement for travel, accommodation, and conference fees to speak at events. J.D.E. is a co-founder of TEGA Therapeutics. J.D.E. and The Regents of the University of California have licensed a University invention to and have an equity interest in TEGA Therapeutics. The terms of this arrangement have been reviewed and approved by the University of California, San Diego in accordance with its conflict-of-interest policies. J.L. is a founder of Glycan Therapeutics and holds equity. G.S. is an employee of Glycan Therapeutics and holds equity. The other authors declare no competing interests.

## Supplemental Figure Legends

**Figure S1.**
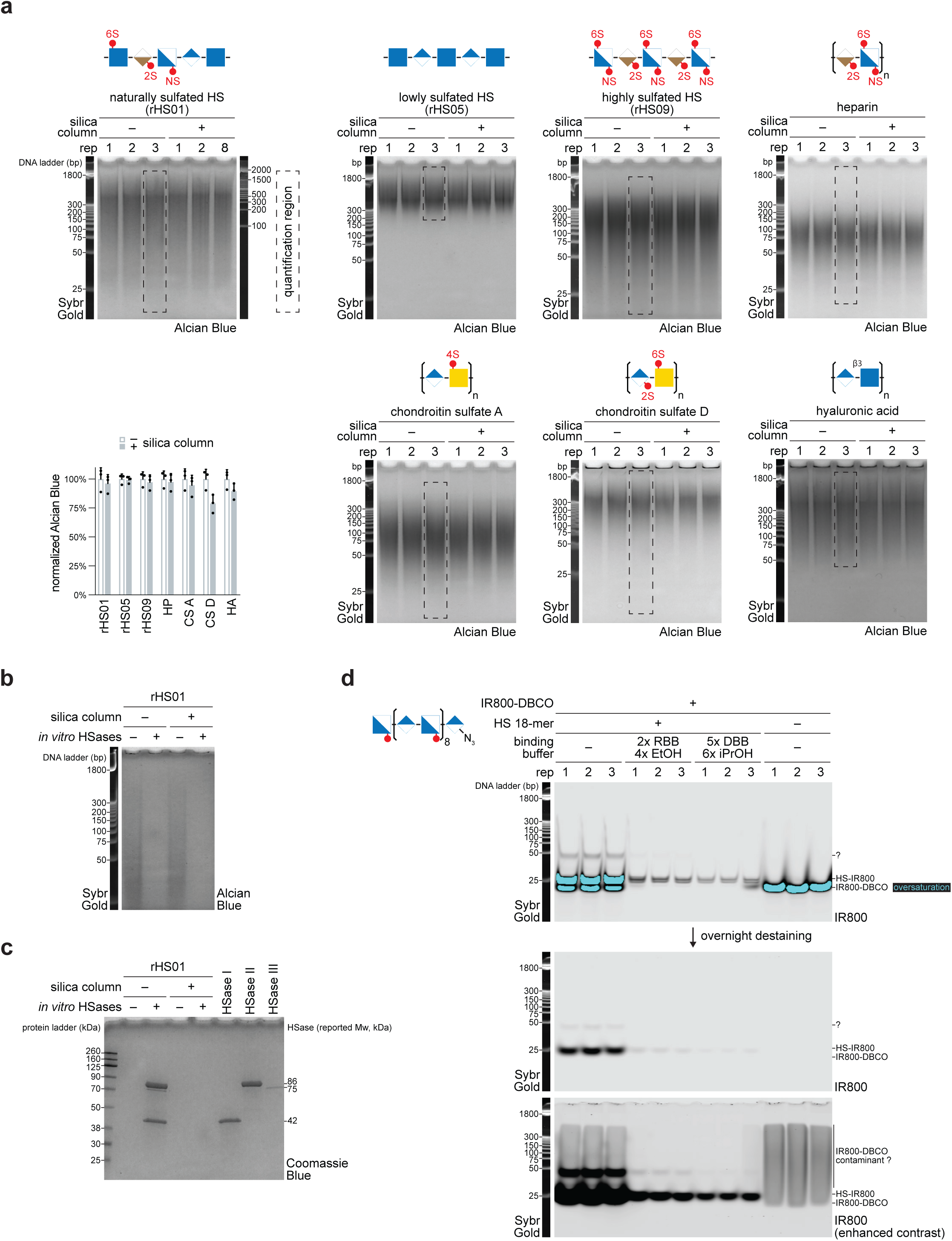
Heparan sulfate can be purified on a silica column. **a.** Cell-derived heparan sulfate was analyzed by native PAGE either directly or following purification on a silica column. HS species rHS01, rHS05, and rHS09 were purified on silica, eluted, and resolved by native PAGE alongside a SYBR Gold-stained DNA ladder. Gels were stained with Alcian Blue, imaged by brightfield imaging, and quantified using ImageJ. Right, quantification of Alcian Blue signal following silica column purification, normalized to the directly loaded (no purification) condition (mean ± s.d., *n* = 3). **b.** Native PAGE analysis of heparan sulfate with or without *in vitro* digestion by heparin lyases I, II, and III, and with or without silica column purification. **c.** SDS-PAGE analysis of heparin lyases I, II, and III under the conditions used in **b**, comparing directly loaded samples with samples subjected to silica column purification. Proteins were resolved alongside a Chameleon Duo protein ladder, stained with Coomassie Blue, and imaged on a Bio-Rad imaging system. Heparin lyases I, II, and III were loaded as controls. **d.** Fluorescent labeling and silica column purification of a synthetic azide-modified HS 18-mer (GlcNS-GlcA repeat). The oligosaccharide was labeled by copper-free click chemistry with IR800-DBCO, split into triplicates, and either purified under different silica column binding conditions or loaded without purification. An equivalent amount of free IR800-DBCO was loaded as a control. Samples were resolved on a 20% native PAGE gel and imaged immediately or after overnight destaining in water.

**Figure S2.**
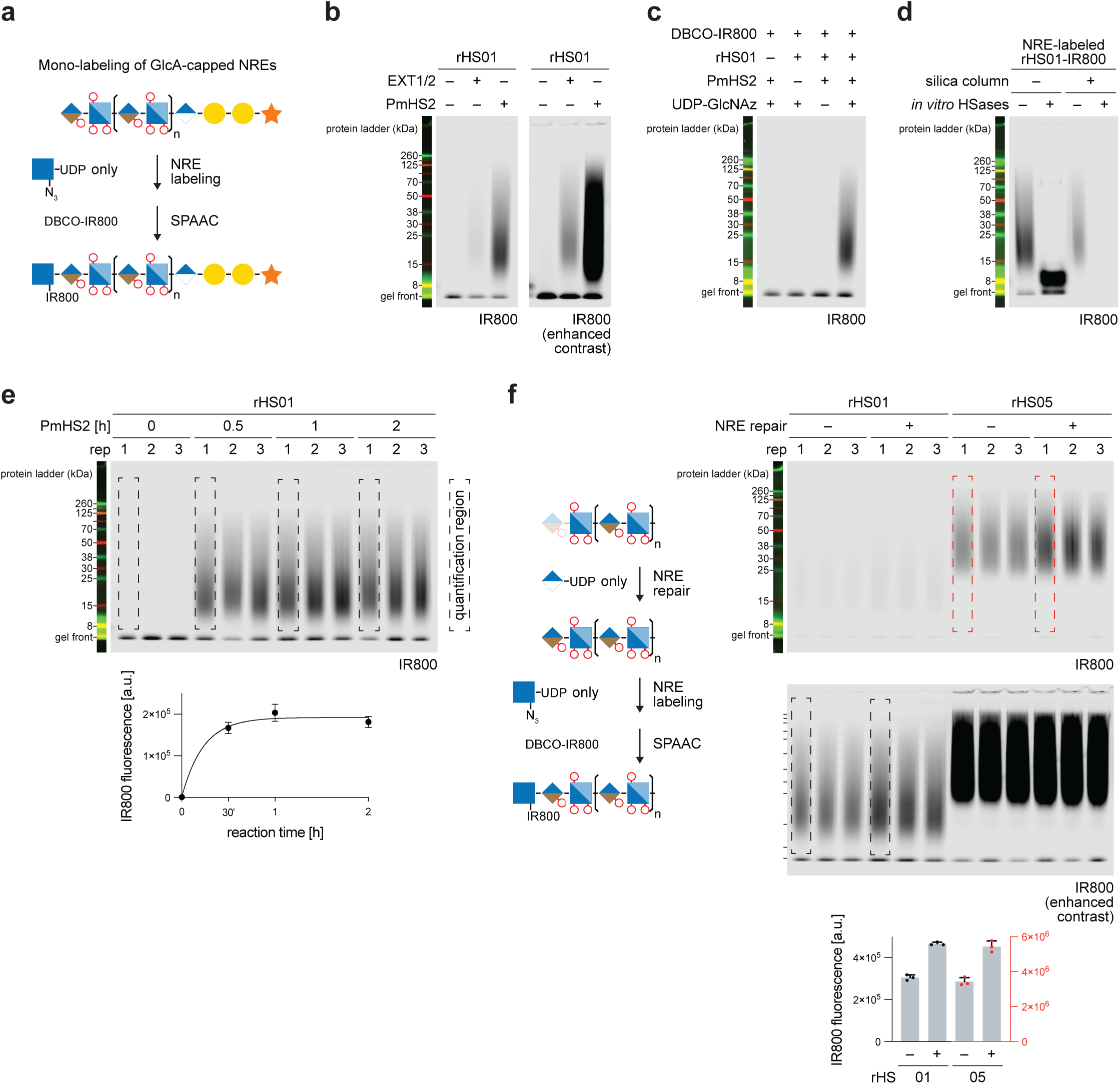
Heparan sulfate can be chemoenzymatically mono-labeled at the non-reducing end. **a.** Schematic of chemoenzymatic mono-labeling of GlcA-capped non-reducing ends (NREs) on native heparan sulfate chains using UDP-GlcNAz only, followed by copper-free click (SPAAC) with IR800-DBCO. **b.** SDS-PAGE analysis of cell-derived heparan sulfate (rHS01) labeled with UDP-GlcNAz by either PmHS2 or EXT1/2. Reactions were incubated for 1 hour using different enzyme concentrations (see Methods). The gel was resolved by SDS-PAGE and imaged using infrared fluorescence detection. **c.** Dependence of mono-NRE labeling on the presence of heparan sulfate, PmHS2, and UDP-GlcNAz. The gel was resolved and imaged as in **b**. **d.** Mono-NRE-labeled heparan sulfate with or without *in vitro* digestion by heparin lyases I, II, and III, and with or without silica column purification. The gel was resolved and imaged as in **b**. **e.** Time-course of PmHS2-mediated mono-NRE labeling with UDP-GlcNAz only, with IR800 fluorescence quantified from the indicated gel region. The gel was resolved and imaged as in **b**. Data were fitted to a one-phase exponential decay model using GraphPad Prism (mean ± s.d., *n* = 3). The negative control lacked PmHS2 but included all other reaction components. **f.** Mono-NRE-labeling using UDP-GlcA only extension (“NRE repair”), followed by UDP-GlcNAz-only labeling of naturally sulfated (rHS01) and minimally sulfated (rHS05) heparan sulfate. The gel was resolved and imaged as in **b**.

**Figure S3.**
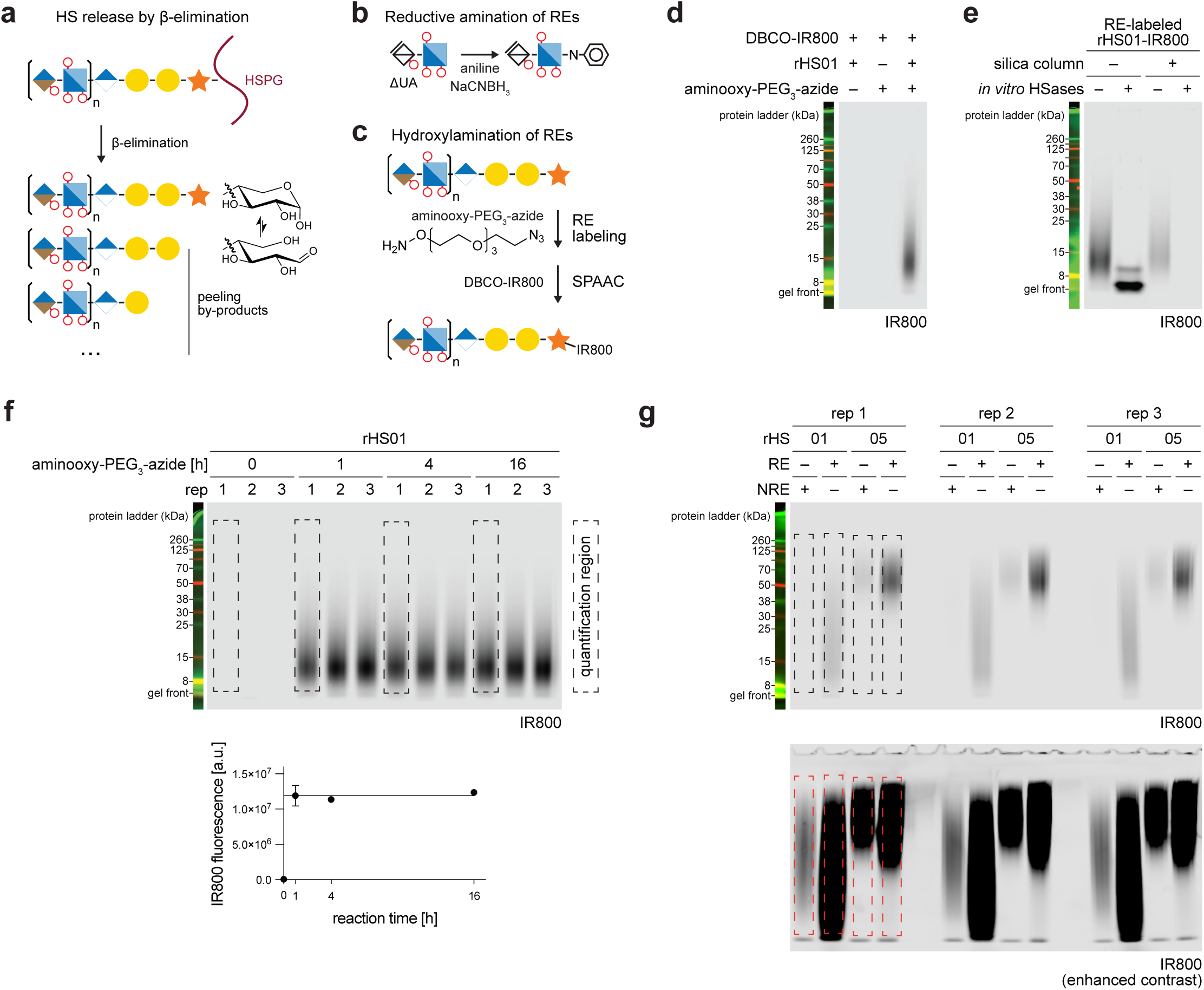
Heparan sulfate can be hydroxylaminated at the reducing end. **a.** Schematic of heparan sulfate release from proteoglycans by β-elimination. The reaction generates peeling by-products, each retaining an aldehyde-reactive hemiacetal at the reducing end. **b.** Schematic of reductive amination of HSases-generated disaccharides using aniline. **c.** Schematic of hydroxylamination of intact heparan sulfate at the reducing end using aminooxy-PEG_3_-azide, followed by copper-free click (SPAAC) with IR800-DBCO. **d.** Dependence of reducing end (RE) labeling on the presence of heparan sulfate and aminooxy-PEG_3_-azide. The gel was resolved by SDS-PAGE and imaged using infrared fluorescence detection. **e.** RE-labeled heparan sulfate with or without *in vitro* digestion by heparin lyases I, II, and III, and with or without silica column purification. The gel was resolved and imaged as in **d**. **f.** Time-course of RE labeling with aminooxy-PEG_3_-azide, with IR800 fluorescence quantified from the indicated gel region. The gel was resolved and imaged as in **d**. Data were fitted to a one-phase exponential decay model using GraphPad Prism (mean ± s.d., *n* = 3). The negative control lacked aminooxy-PEG_3_-azide but included all other reaction components. **g.** Comparison of NRE labeling (following NRE repair) versus RE labeling of naturally sulfated (rHS01) and minimally sulfated (rHS05) heparan sulfate. IR800 fluorescence was quantified from the indicated gel region. The gel was resolved and imaged as in **d**.

**Figure S4.**
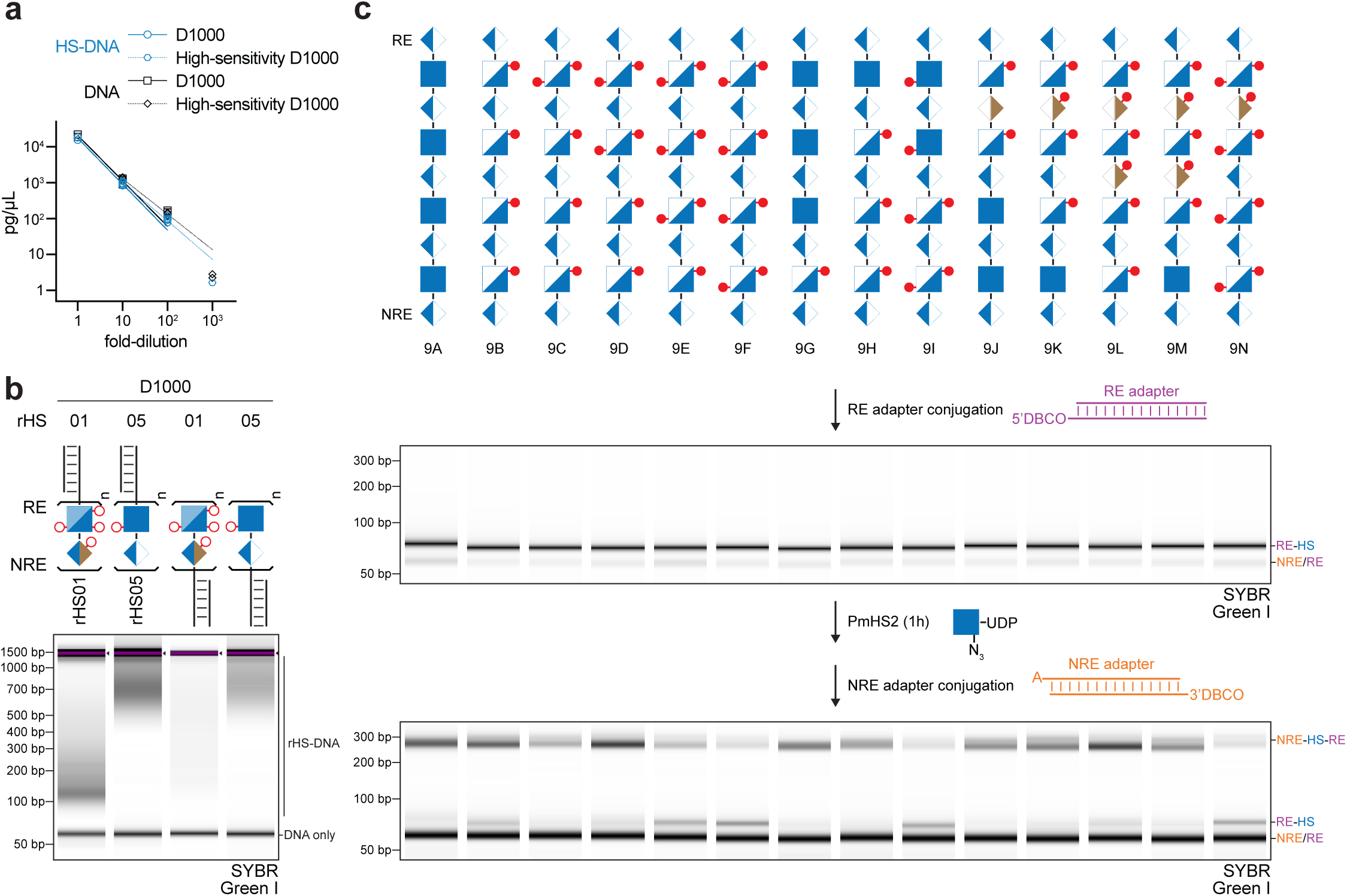
TapeStation analysis of HS-DNA and DNA-HS-DNA chimeras. **a.** Quantification of HS-DNA and DNA detection sensitivity across a dilution series by D1000 and High-Sensitivity D1000 TapeStation assays. Curve represents a nonlinear regression fit to a log-log line model, with both axes displayed on logarithmic scales (GraphPad Prism). Individual data points shown; n = 3 except at the highest dilution, where n = 1 or 2, as indicated in the figure. **b.** TapeStation (D1000) analysis of cell-derived rHS01 and rHS05 conjugated to a DNA adapter at either the RE or NRE. **c.** TapeStation (D1000) analysis of DNA-HS-DNA chimeras assembled from 14 synthetic HS 9-mers of varying sulfation. Each synthetic HS 9-mer was conjugated to a DNA adapter at the RE end, labeled at the NRE for 1 h, then conjugated to a second adapter at the NRE.

**Figure S5.**
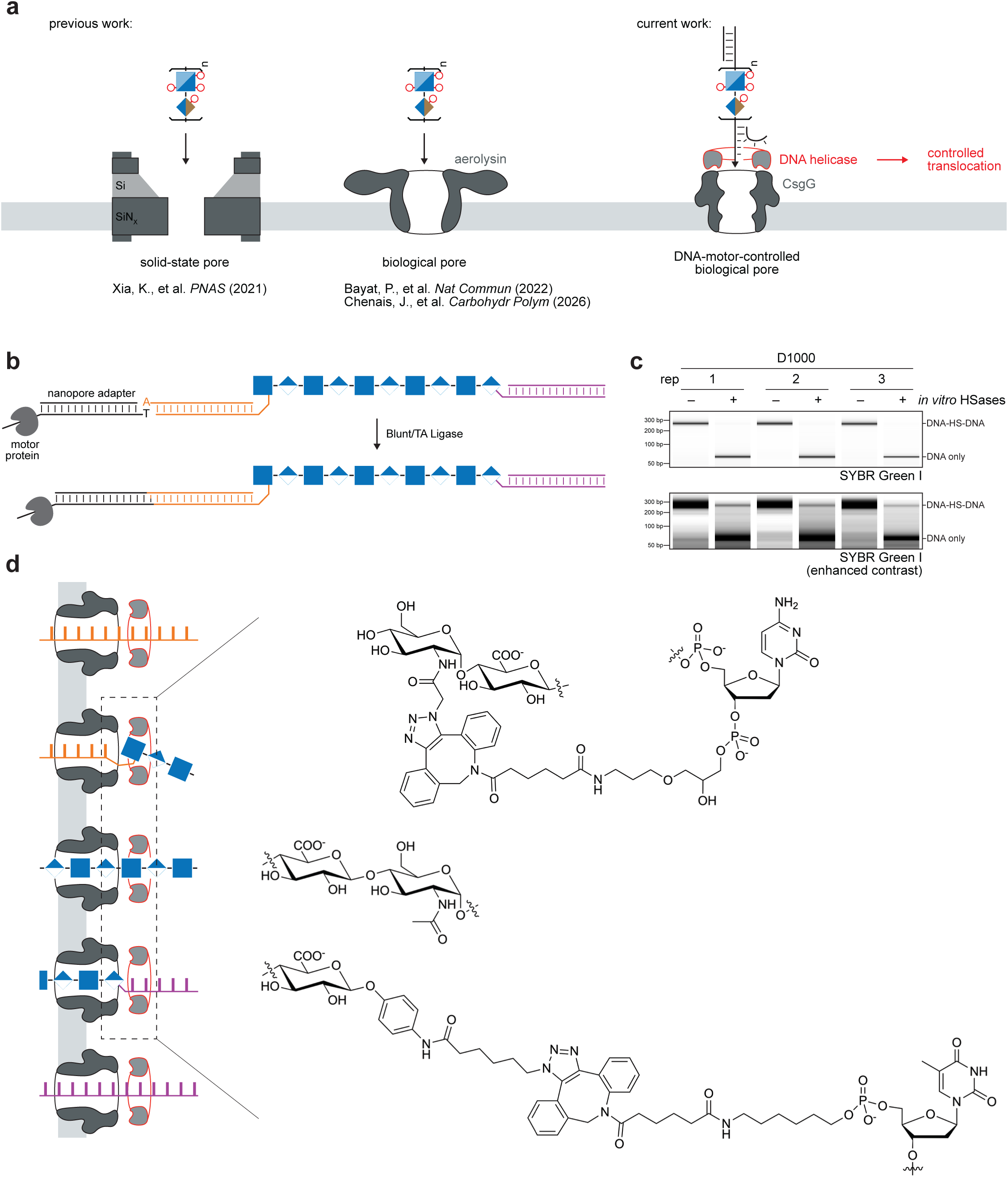
Framework for nanopore analysis of HS. **a.** Comparison of nanopore approaches for glycosaminoglycan (GAGs) analysis. Previous approaches used solid-state pores^20^ or biological aerolysin pores^19,21^. This work uses a commercial DNA-motor-controlled CsgG biological pore, in which a DNA helicase ratchets the DNA-HS-DNA chimera through the pore for controlled translocation. **b.** Schematic of nanopore adapter ligation to a DNA-HS-DNA chimera via Blunt/TA Ligase. The motor protein-containing adapter is attached to the NRE. **c.** TapeStation (D1000) analysis of DNA-HS-DNA chimeras with (+) and without (-) HSases treatment. n = 3 technical replicates. **d.** Chemical structures of the NRE (top) and RE (bottom) HS-DNA junctions.

**Figure S6.**
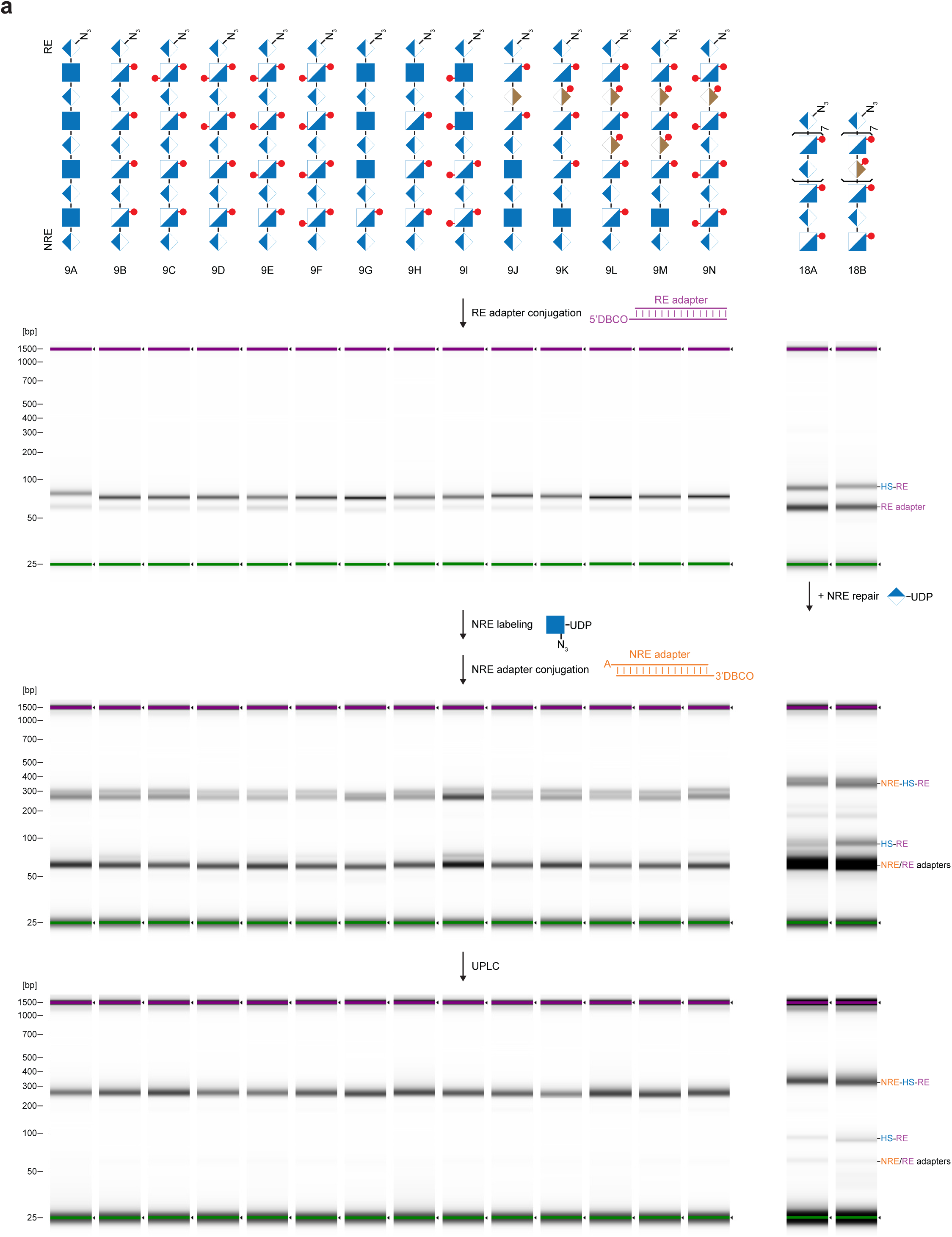
Conjugation and purification of synthetic HS standards. **a.** TapeStation (D1000) analysis of DNA-HS-DNA chimeras assembled from 14 synthetic HS 9-mers and 2 18-mers of varying sulfation. Each synthetic HS standard was conjugated to a DNA adapter at the RE end, labeled at the NRE for 4 h, conjugated to a second adapter at the NRE, and purified by UPLC. SYBR electropherograms are uniformly normalized across the experiment.

**Figure S7.**
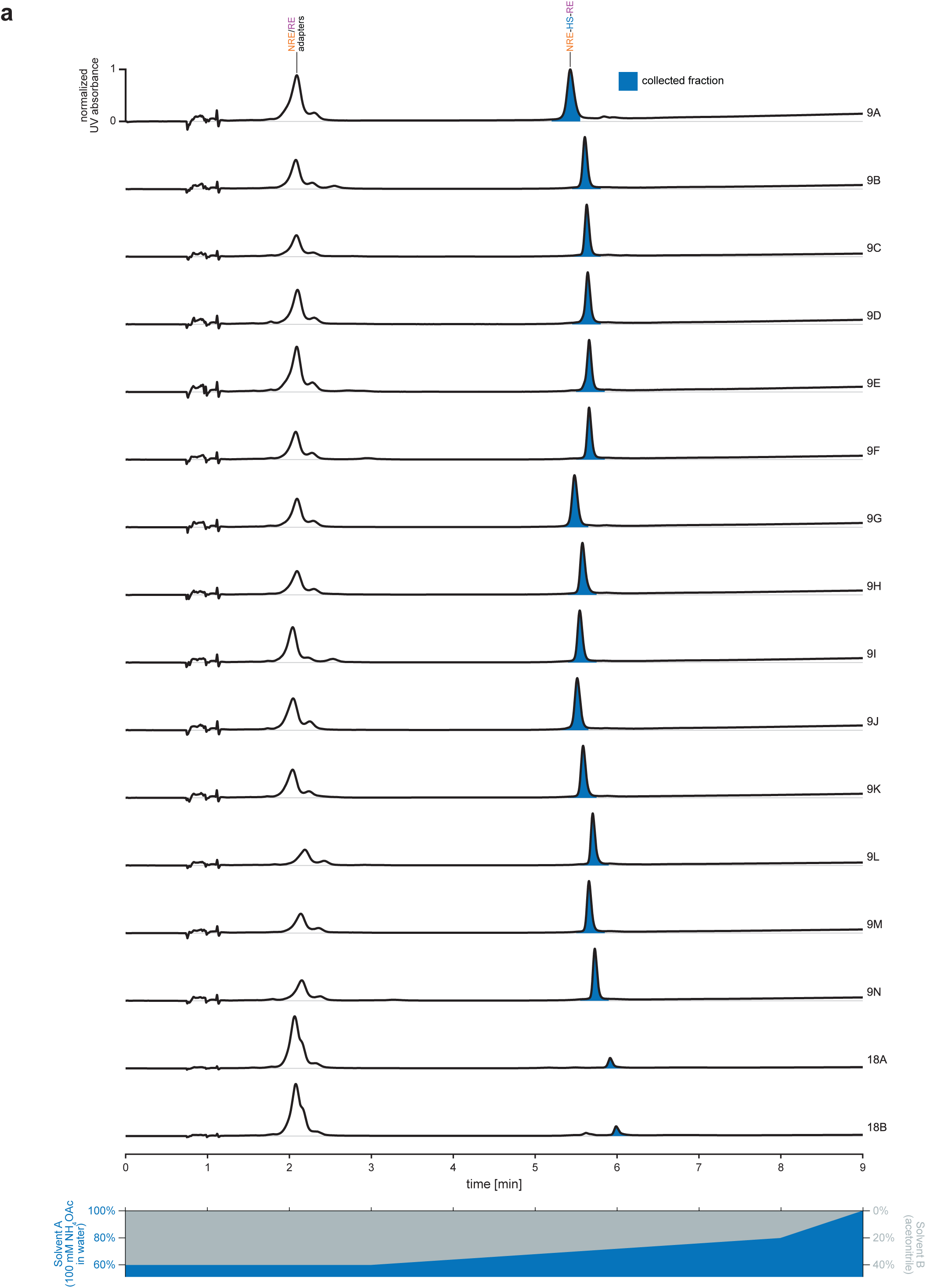
UPLC purification of DNA-HS-DNA chimeras. **a.** Representative UPLC UV absorbance traces from purification of DNA-HS-DNA chimeras from various synthetic standards. Annotated peaks correspond to the DNA adapter, HS-DNA intermediate, and full DNA-HS-DNA chimera. The collected fraction is highlighted in blue.

**Figure S8.**
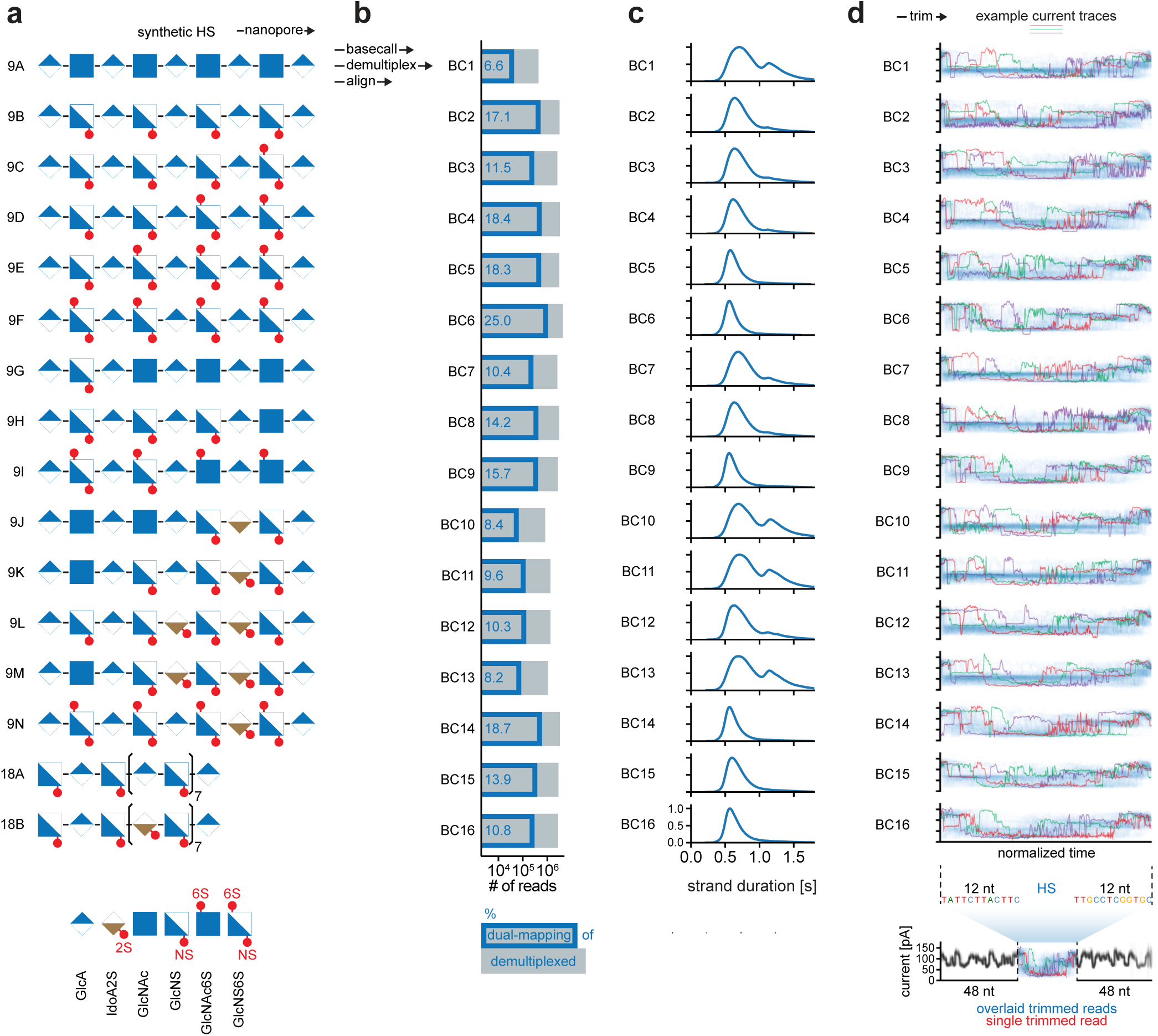
Additional characterization related to Figure 4. **a.** Panel of synthetic HS standards for nanopore current fingerprinting. SNFG structures are shown for 16 standards, comprising fourteen 9-mers and two 18-mers, with sulfation positions annotated. Bottom, SNFG legend for HS monosaccharide building blocks with sulfation position annotations (2S, NS, 6S). **b.** Bar plot of total demultiplexed reads overlaid with dual-mapping reads. Barcoded DNA-HS-DNA chimeras were nanopore-sequenced, base-called, demultiplexed, and aligned to an adapter reference containing the NRE and RE adapters. **c.** Strand-duration distributions of dual-mapping DNA-HS-DNA reads shown as kernel density estimates. **d.** Overlaid nanopore current traces for each HS standard. HS-associated current segments were trimmed with 12 nt of flanking DNA adapter sequence on each side. Blue, overlay of 100 reads per standard; red, green, and purple - representative single reads. Bottom, full-length trace for 18B indicating the trimmed region used for comparison.

**Figure S9.**
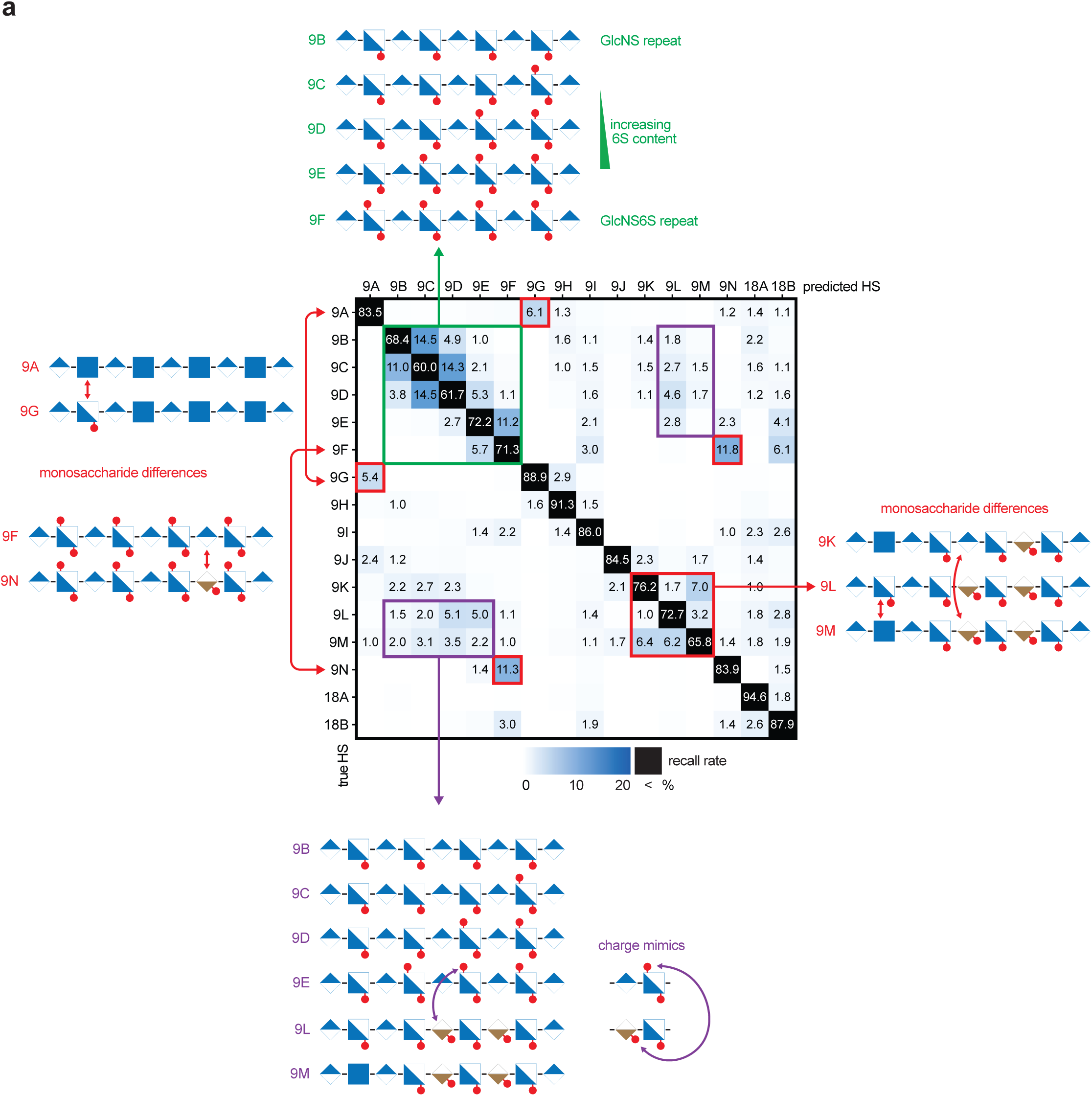
Misclassification patterns across the confusion matrix. **a.** Confusion matrix showing per-class recall (%) across the 16-standard HS panel. Recall rates higher than 1% are shown on a color gradient, with rates higher than 20% depicted in black. Clusters of misclassifications are highlighted by color. Red, pairs differing by a single monosaccharide (9A/9G, 9F/9N, 9K/9M, and 9L/9M). Green, GlcNS repeat series with increasing 6S content grading toward the GlcNS6S repeat. Purple, GlcA-GlcNS6S and IdoA2S-GlcNS motifs as charge mimics.

**Figure S10.**
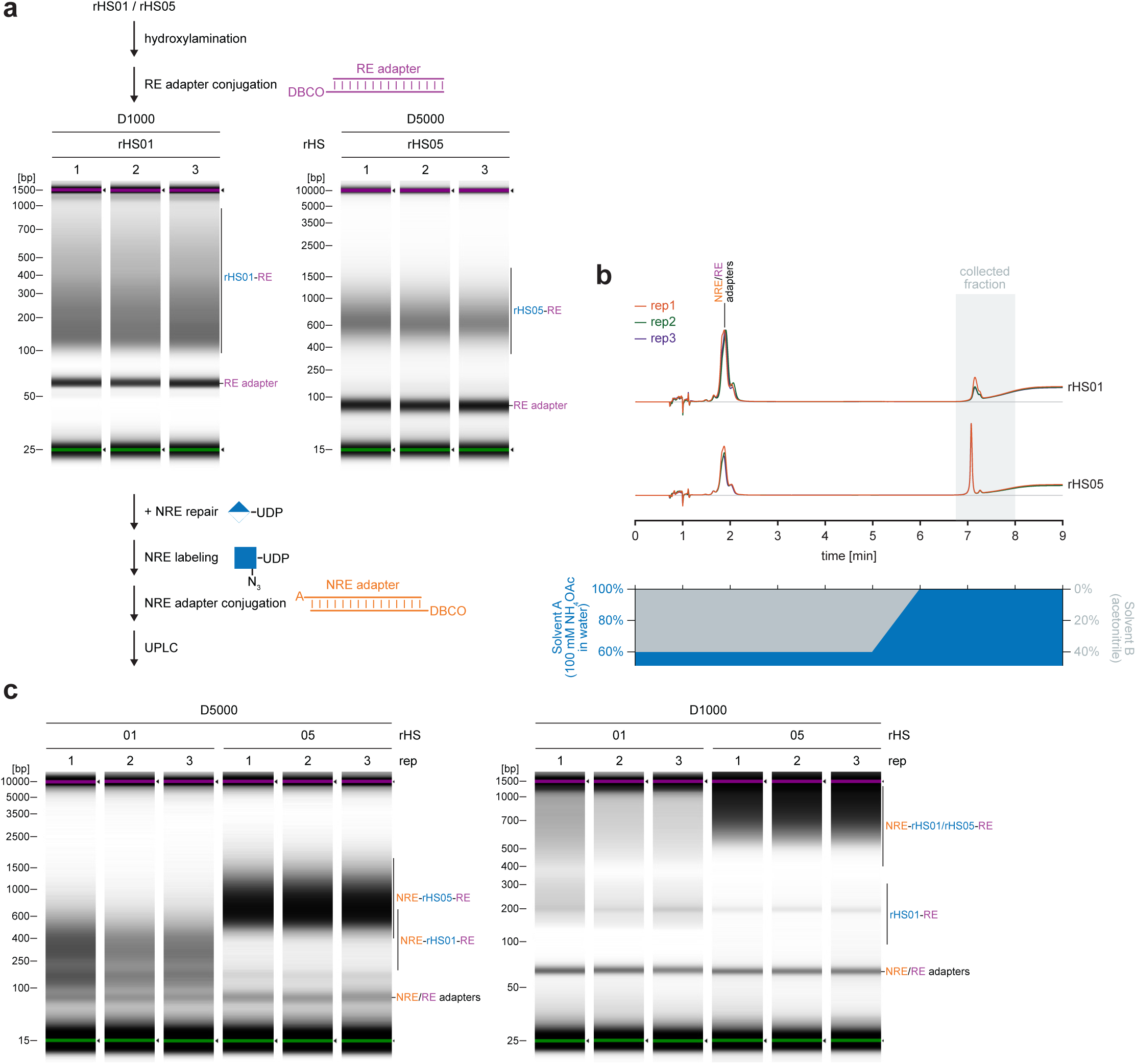
Cloning and purification of cell-derived HS standards. **a.** TapeStation analysis of DNA-HS-DNA chimeras assembled from 2 cell-derived rHS of varying sulfation. Each cell-derived HS was labeled at the reducing end (RE), conjugated to a DNA adapter at the RE, NRE-repaired and labeled at the non-reducing end (NRE), conjugated to a second adapter at the NRE, and purified by UPLC. SYBR electropherograms are uniformly normalized across the experiment. **b.** Representative UPLC UV absorbance traces from purification of DNA-rHS-DNA chimeras from cell-derived HS. Annotated peaks correspond to the DNA adapter and full DNA-HS-DNA chimera. The collected fraction is highlighted in blue. **c.** TapeStation of the UPLC purified material used as input in the pore. SYBR electropherograms are uniformly normalized across the experiment.

## Supplemental Tables

**Table S1. Nanopore read statistics.**

**Table S2. Heparan sulfate chains.**

**Table S3. DNA oligos.**

